# Spatiotemporally regulated transcriptional networks control the plasticity of chickpea root exodermis

**DOI:** 10.1101/2025.01.03.631192

**Authors:** Leonardo Jo, Rianne Kluck, Sara Buti, Julia Holbein, Martin Rázus, Virevol Thakro, Job Naaijer, Himanshu Chhillar, Pingtao Ding, Rochus Benni Franke, Kaisa Kajala

## Abstract

Abscisic acid (ABA) regulates plant responses to stress and influences the differentiation of root barrier cell types, such as the endodermis and exodermis. Despite the importance of the exodermis in limiting water and solute fluxes, its regulation remains poorly understood in legumes. Here, we characterize suberization and lignification induced by exogenous ABA in tissues across eight galegoid legumes to identify exodermis-forming species in the clade. Chickpea deposited suberin lamella specifically in the outermost cortex and formed a functional apoplastic barrier, i.e. an exodermis. Transcriptomics of chickpea roots treated with ABA revealed ABA-induced programs that were temporally separated, specifically, a rapid program with the general ABA response, and a delayed programs of suberin and lignin biosynthesis. We identified WRKY and MYB transcription factors that putatively link the ABA response to the suberin biosynthesis, and using single-cell RNA-seq of chickpea roots we inferred an exodermis-expressed WRKY-MYB regulatory unit upstream of the suberin biosynthetic genes. Transactivation assays supported an ABA-dependent transcription factor activity upstream of suberin biosynthesis pathway in chickpea. Our results reveal a cell-type specific transcription factor hierarchy that coordinates hormone perception into an important mechanism of root plasticity in legumes.

## Introduction

Plants exhibit indeterminate and highly plastic growth, allowing them to adjust their development in response to fluctuating environmental conditions. This plasticity extends to the differentiation trajectories of individual plant cells, which are not solely determined by developmental programs but also integrate environmental signals to modify their structure and function. One of the key stress cues in plants is abscisic acid (ABA), a plant hormone that regulates many responses to drought, salinity, and other abiotic stressors (Muhammad Aslam et al. 2022). ABA has been shown to influence the differentiation of various root cell types, including the endodermis and the exodermis. In these barrier cell types, ABA induces a differentiation program that deposits suberin into the cell wall (Barberon et al. 2016; Shukla et al. 2021; Shiono et al. 2022; Cantó-Pastor et al. 2024; Chen et al. 2024). Suberin deposition significantly contributes to the apoplastic diffusion barrier properties of these cell types and helps with retention of water and likely oxygen (Shiono et al. 2022; Su et al. 2023; Cantó-Pastor et al. 2024). Notably, endodermis and exodermis also have another differentiation program that deposits localized lignin into their cell walls, and this program is in many species under developmental control (Alassimone et al. 2010; Roppolo et al. 2011; Naseer et al. 2012; Hosmani et al. 2013; Lee et al. 2013; Doblas et al. 2017; Nakayama et al. 2017; Manzano et al. 2024). The endodermal lignin is precisely positioned and forms the Casparian strip that divides the endodermal cells to soil-facing and vasculature-facing sides and functions as an apoplastic diffusion barrier between the two. The Casparian strip deposition machinery is controlled by a MYB (MYELOBLASTOSIS) family transcription factor (TF) MYB36 (Kamiya et al. 2015; Liberman et al. 2015), and it is precisely localized via the Schengen pathway (Alassimone et al. 2010; Roppolo et al. 2011; Hosmani et al. 2013; Lee et al. 2013; Doblas et al. 2017; Nakayama et al. 2017). This provides the endodermis the ability to be a selective control point for two-way fluxes between the root and the soil (Barberon 2017). In contrast, the exodermal lignification patterns include outward-facing polar cap (Manzano et al. 2024) and non-polar deposition (Jones et al. 2025), and both the exodermal lignin (transcriptional) regulators and players involved in the positioning remain largely unknown. Together, the endodermis and the exodermis coordinate their critical roles in regulating the two-way fluxes of water, solutes and gases and protecting plants against environmental stressors. At the same time, they each integrate numerous environmental cues into their differentiation programs to optimize their functions.

While the endodermis is one of the best characterized plant cell types for its development and differentiation (Geldner 2013; Andersen and Drapek 2024), the exodermis remains relatively unexplored because the model for root development, *Arabidopsis thaliana*, lacks this cell type (Liu and Kreszies 2023). However, the exodermis in its diverse morphologies is found in around 90% of flowering plants (Perumalla et al. 1990b, 1990a; Peterson and Perumalla 1990). Considering the ABA-induced differentiation program in endodermis and exodermis, there is diversity also in its regulation across the flowering plants but universally, the suberin deposition program is regulated by a group of closely related MYB TFs (Lashbrooke et al. 2016; Baumgart et al. 2025). In Arabidopsis, ABA induces the deposition of endodermal suberin lamella (SL) via multiple, redundantly acting MYBs from clades S10, S11 and S24 (Dubos et al. 2010; Du et al. 2015) including AtMYB39, AtMYB41, AtMYB53, AtMYB92 and AtMYB93 (Cohen et al. 2020; Shukla et al. 2021; Xu et al. 2022). Similarly in rice, four MYBs, OsMYB39a, OsMYB41, OsMYB92a, and OsMYB92b, were identified as the redundant core regulators of endodermal suberin, which also were responsive to ABA (Chen et al. 2024). Intriguingly, these rice MYB TFs do not affect the exodermal suberin which also are deposited in rice roots in response to ABA, suggesting that a separate set or additional TFs are required for exodermal suberin in rice. In contrast, tomato has suberin only in the exodermis and the homologs of these MYBs, specifically SlMYB92, SlMYB41 and SlMYB63, regulate the exodermal SL in a non-redundant fashion (Cantó-Pastor et al. 2024; Jo et al. 2025). So, a set of conserved MYBs regulates the suberin differentiation program, which is ABA-responsive in endodermis, exodermis, or both, depending on the angiosperm species. In addition to MYBs, suberin biosynthesis is also regulated by some of the WRKY family TFs. Specifically, AtWRKY9 (Krishnamurthy et al. 2021) and AtWRKY33 (Krishnamurthy et al. 2020) directly induce expression of Cytochrome P450 genes (*CYPs*) encoding AtCYP94s and AtCYP86s that carry out fatty acid ω-oxidation in suberin biosynthesis (Serra and Geldner 2022). In tomato, SlWRKY71 was shown act antagonistically to SlMYB93 and SlMYB41, and repress expression of suberin biosynthesis genes (Jo et al. 2025). Ultimately, the direct link between ABA signalling and these cell-type-specific differentiation programs coordinated by the MYBs and WRKYs remains unknown.

Exodermis has been reported to be found in the majority of angiosperm families, with only a small number of families having species either with and without an exodermis (Perumalla et al. 1990b, 1990a; Peterson and Perumalla 1990; Artur and Kajala 2021). Legumes (Fabaceae) were originally reported as one of the families not containing species with an exodermis (Perumalla et al. 1990b) and it was hypothesized that the presence of a barrier at the outermost cortex layer would be inhibitory to the legumes’ ability to form symbioses with nitrogen-fixing bacteria in root nodules formed from the cortex layer (Enstone et al. 2002). In the past two decades there has been contrasting reports of the presence of exodermis in different legumes such as pea (*Lathyrus oleraceus*, previously *Pisum sativum*) (Perumalla et al. 1990b; Taleisnik 1999; Bani et al. 2018) and chickpea (*Cicer arietinum*) (Schreiber et al. 1999; Sharma et al. 2023), and this may be due to the stress-induced, dynamic nature of the exodermis in many species. Intrigued by this gap of knowledge and potential diversity of root barriers in legumes, we set out to characterize the ABA-responsive differentiation programs in a selection of galegoid legumes (Fig. 1a) such as chickpea, lentil (*Lens culinaris*) and alfalfa (*Medicago sativa*) to query how exodermis is regulated in this clade. Our results provide key insights into the diversity of cell-type specific responses to ABA across legume species and identify core components of the regulatory network that translate hormone perception into exodermis differentiation in chickpea.

**Figure 1.**
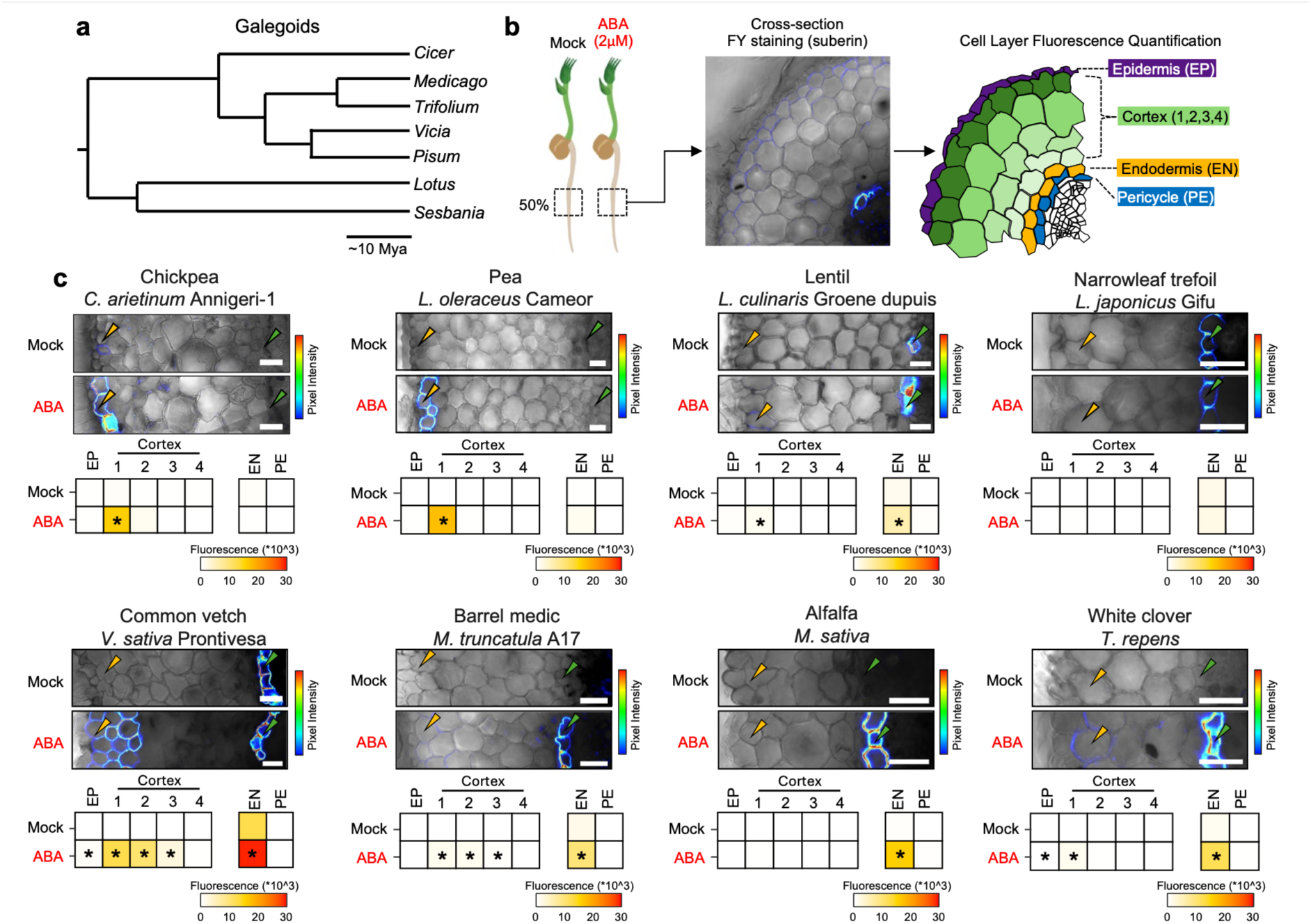
Galegoid legumes deposit suberin to the cell walls of different root cell types in response to ABA. **a,** Representative phylogenetic tree of the galegoid legume clade. The tree was adapted from (Cannon et al. 2010). *Lens culinaris* is nested in the genus *Vicia* (Ellis et al. 2024). **b,** Graphical representation of the survey for observing cell-type suberization in legume roots treated with mock or ABA (2 µM) for 3 days. The middle point of the root (50%) was cross-sectioned, stained with FY and counter-stained with aniline blue. Finally, the fluorescence levels of cell walls of individual cell layers of the root epidermis (EP), the four outermost cortex layers (C1 to C4) out of the up to 12 cortex layers (depending on the species), the endodermis (EN) and the pericycle (PE) were quantified. **c,** Quantification summary of FY intensity across multiple cell layers of legume roots in response to ABA. For each species, the top images show representative confocal images of cross-section of roots treated with mock or ABA, with the FY signal overlaid on top of bright field images (scale = 25 µm). The yellow and green arrow depicts the position of the outermost layer of the cortex (C1) and endodermis, respectively. On the bottom, the heatmaps show the average pixel intensity of the FY signal in different layers of the root cross section. Asterisks indicate significance based on Wilcoxon rank-sum test with a Benjamini-Hochberg adjusted P-value. Threshold of 0.05 was used to indicate the statistical significance (n ³ 8).

## Materals and methods

### Sterilization and germination of legume seeds

Legume seeds were obtained from the following sources: *C. arietinum* Annigeri-1 accession ICC 5679 from ICRISAT; *L. oleraceus* Cameor (Kreplak et al. 2019) from Judith Boursin, *L. culinaris* Groene Dupuis and *V. sativa* Prontivesa from Vreeken’s Zaden (the Netherlands), *L. japonicus* Gifu from Tonni Andersen, *M. truncatula* A17 from Wouter Kohlen, and *M. sativa* and *T. repens* from Intratuin B.V. (the Netherlands).

Seeds from *C. arietinum* Annigeri-1, *L. oleraceus* Cameor, *L. culinaris* Groene Dupuis, *V. sativa* Prontivesa, *T. repens,* and *M. sativa* were gas sterilized by placing the seeds in a desiccator with a mixture of 50 mL of commercial bleach and 2.5 mL of HCl for 12 hours. After sterilization, seeds were soaked in distilled water for 12 hours in the dark at room temperature. Soaked seeds were placed onto sterile 0.7% (w/v) Phyto agar (Duchefa) dissolved in water. Seeds were then germinated in the dark for 3 days at 25°C. For the germination, plates were kept in a 60° to 70° angle to ensure a uniform growth direction of the roots. For *M. truncatula* A17, seeds were scarified for 7 minutes using sulfuric acid and sterilized for 7 minutes with 10% (v/v) commercial bleach. After sterilization, the seeds were washed five times with distilled water and kept in water for 12 hours. After soaking, seeds were placed on petri dishes containing 1% (w/v) Phyto agar and cold stratified in the dark at 4°C for 3 days. To initiate germination, the petri dishes were inverted horizontally (with the medium on top) and transferred to growth cabinets at 25°C in the dark for 12 hours. Germinated seedlings were then transferred to 0.7% (w/v) Phyto agar plates and kept in the dark for another 36 hours in a 60° to 70° standing angle. For *L. japonicus* Gifu, seeds were mechanically scarified with sandpaper and surface sterilized with 10% commercial bleach for 10 minutes. After sterilization, seeds were washed five times and soaked in water at room temperature for 12 hours. Seeds were then transferred to plates with 1% (w/v) Phyto agar and cold-stratified at 4°C in darkness for 3 days. Seeds were germinated at 25°C in darkness for five days and plates were placed in 60-70° angle.

The growth conditions for each experimental approach are described within each section, but in general, the 3-day treatments were carried on plates where root growth can be monitored, while 24-h or shorter treatments were carried on liquid media in magenta boxes to ensure ABA exposure to all root surfaces.

### Suberin and lignin histology and fluorescent signal quantification

Four-day old seedlings of *C. arietinum* Annigeri-1, *P. sativum* Cameor, *L. culinaris* Groene Dupuis, *V. sativa* Prontivesa, *T. repens* and *M. sativa*, two-day old seedlings from *M. truncatula* A17, and five-day old seedlings from *L. japonicus* Gifu were transferred to ½ MS square plates supplemented with 2 µM ABA or mock conditions. The seedlings were maintained in a D-root system (Silva-Navas et al. 2015) in growth chamber under long-day conditions (16 h light, 8 h dark; 25°C; 70% humidity). After 3 days, 1-cm root segments at the middle point of the root (50% region) were vacuum-infiltrated for 1 h in a solution containing 4% (w/v) paraformaldehyde (PFA) dissolved in PBS buffer. After vacuum infiltration, roots were kept in 4% PFA for at least 12 hours at 4°C and subsequently washed three times with PBS. Root segments were embedded in 4% (w/v) agarose (*C. arietinum*, *P. sativum*, *L. culinaris*, *V. sativa*) or 3% (w/v) (*M. truncatula*, *M. sativa*, *L. japonicus*, *T. repens*) and cross-sectioned into 250 µm thick sections using the VT1000 S vibratome (Leica). Fluorescent staining and confocal imaging of suberin with FY and lignin with BF in root cross sections was performed as follows (Jo et al. 2025). For suberin visualization, root cross-sections were stained with 0.01% (w/v) FY 088 dissolved in 96% EtOH under continuous agitation for 20 min in dark at RT. Then, to counterstain all other cell structures, the FY was replaced with 0.5% (w/v) aniline blue in Milli-Q water for another 20 min. The sections were then washed twice with Milli-Q until the excess water ran clear and kept in 50% (v/v) glycerol until imaging. For lignin visualization, root cross-sections were stained with 0.02% (w/v) BF dissolved in ClearSee under continuous agitation for 20 min in dark at RT. The root cross-sections were washed four times with ClearSee in intervals of 15 minutes with agitation in the dark at RT.

The quantification of FY and BF fluorescent signals along individual cell layers of the root were done using the Icy software (https://icy.bioimageanalysis.org/). To analyse fluorescence intensity, 2D polylines were manually traced along the anticlinal cell walls of at least 10 individual cells from different cell layers within each root cross-section. In sections where FY and BF were clearly visible in the exodermis and endodermis, regions with the clearest FY or BF signal was selected for the quantification. Following this reference point, we measured the fluorescent intensity of layers above and below these cells to extract the signal in adjacent cell-types (i.e. epidermis and cortex) hence the results represent overall increase in signal in each cell layer, not increase in the number of cells with signal (which is shown in Supplementary Fig. S3). The ‘Path Intensity Profile’ plugin in Icy was used to measure the maximum pixel intensity for each cell wall in the FY or BF channels. For background correction, random points along the root surface were selected to estimate the overall background signal for each image. The average background signal of each image was subtracted from the maximum pixel intensity of each cell wall. Finally, for each image, the fluorescence signal of the cell layer was determined as average signal intensity across all measured cells of the layer. Quantification was done using cross-sections of at least 8 individual roots in mock or ABA conditions and statistical differences were determined based on Wilcoxon rank-sum test with a Benjamini-Hochberg adjusted P-value threshold of 0.05 was used to indicate the statistical significance.

### GC-MS

For the profiling of suberin monomers, four-day-old seedlings of *C. arietinum* Annigeri-1 and *V. sativa* Prontivesa and two-day-old seedlings of *M. truncatula* A17, were transferred to magenta boxes filled with sterile ½ MS medium supplemented with or without 2 µM ABA. A metal grid was placed in the magenta boxes to ensure that only the root was in direct contact with the liquid medium. The seedlings were maintained in the magenta boxes under long-day conditions (16 h light, 8 h dark; 25°C; 70% humidity), under slow agitation (20 rpm) for 24 hours. Root samples (root tip to the 50% region of the root) were collected for an average of 350, 210 and 50 mgs of fresh weight (FW) tissue per biological replicate for *C. arietinum, V. sativa and M. truncatula*, respectively. Root samples were immediately placed in hot isopropanol (70°C) for 30 minutes. The samples were extracted using a Soxhlet apparatus for 8 hours, first with chloroform followed by methanol, to eliminate all soluble lipids. Subsequently, extracted tissues were dried over silica gel in a desiccator. Suberin monomers were released through transesterification with sulfuric acid in methanol (5% v/v) at 85°C for 3 h. Dotriacontane (0.2 μg/μl) was added to each sample as an internal standard. Sodium chloride (2.5% w/v) and methyl tert-butyl ether (MTBE) were added to stop the transesterification reaction and to extract suberin monomers. The MTBE extract was washed with sodium chloride (0.09% w/v) in Tris buffer (100 mM) at pH 8. The MTBE fraction was then completely evaporated and subsequently derivatized with N,N-bis (trimethylsilyl)trifluoroacetamide (BSTFA) and pyridine at 70°C for 40 minutes. Gas chromatography (GC) equipped with a flame ionization detector (6890 N Network GC System, Agilent Technologies) was used to separate and quantify the compounds. For compound identification, gas chromatography coupled with a mass spectrometry selective detector (GC-MS; 5977A MSD, Agilent Technologies) was utilized and fragmentation spectra were matched against an internal library of known suberin monomers and the NIST database for identification. The method was adapted from (Delude et al. 2017). Full data is given in Supplementary Table S1.

### Transmission electron microscopy (TEM)

Germinated seedlings of chickpea, *M. truncatula* and *V. sativa* were transferred for 3 days on plates with either mock or 2 µM ABA treatment. After three days of treatment, 1 cm root pieces were harvested in the 75% region of the individual roots. The older developmental zone was selected to increase the likelihood of capturing suberized cells (Supplementary Fig. S2). Root pieces were fixed in 2.5% glutaraldehyde + 2% PFA in phosphate buffer (PB 0.1 M) at 4°C for at least two weeks. After incubation, samples were washed 6 times for 10 min in PB, incubated in 1% osmium tetroxide dissolved in PB for 1 h, and washed again 3 times for 10 min using MilliQ water. Samples were dehydrated using a concentration gradient of acetone solutions (10% for 15 min, 30% for 15 min, 50% for 15 min, 70% for 15 min, 80% for 15 min, 90% for 15 min, 96% for 10 min, 100% for 15 min, 100% for 30 min). This was followed by infiltration with LR white in a concentration gradient (1:2 LR white: EtOH for 1 h, 1:1 LR: EtOH for 1 h, 2:1 LR white: EtOH for 1 h, 100% LR white for 2 h, 100% LR white overnight at room temperature, 100% resin for 2 h) and polymerized for 24 h in an oven at 60-65°C. Polymerized samples were cross-sectioned to 50 nm thickness for *M. truncatula* and chickpea, and *V. sativa* was sectioned to 70 nm thickness using a Leica ultramicrotome UC7, before attaching to 100 mesh carbon grids. The grids were incubated for 10 min using 2% uranyl acetate followed by 5 washes with MilliQ water. Next, the grids were incubated with lead citrate (Electron Microscopy Sciences) for 10 min in a CO_2_-free environment. The grids were washed two times with 0.01 N CO_2_-free water followed by 3 final washes with MilliQ water. Sample preparation, processing, and imaging were carried out by the Wageningen Electron Microscopy Centre. A JEOL JEM1400 transmission electron microscope was used to image the samples.

### Propidium iodide diffusion assay

Four-day-old seedlings of *C. arietinum* Annigeri-1 and *V. sativa* Prontivesa and two-day-old seedlings of *M. truncatula* A17, were transferred to new ½ MS square plates supplemented with 2 µM ABA or mock conditions. The seedlings were maintained in a D-root system (Silva-Navas et al. 2015) in growth chamber under long-day conditions (16 h light, 8 h dark; 25°C; 70% humidity). After 3 days, root segments up to the 75% region, including the root tip, were exposed to 30 µg/mL propidium iodide solution in the dark at 28°C for 2 hours. Root segments were washed twice in MilliQ water and 0.5 cm segments from the 50% region of the root were excised and embedded in 4% (w/v) agarose for *C. arietinum* and *V. sativa* and in 3% (w/v) agarose for *M. truncatula*. Root cross-sections (250 µm thick) were obtained using the VT 1000 S vibratome (Leica). Optical laser scanning microscopy was performed on a Zeiss apotome microscope with the ×20 objective at 405 nm excitation and 600–650 nm detection. The quantification of the cell-layer PI intensity was done as described above.

### RNA-seq harvest, libraries, sequencing

Four-day-old seedlings of *C. arietinum* Annigeri-1 were transferred to magenta boxes filled with sterile ½ MS medium supplemented with or without 2 µM ABA. Like the GC-MS experiments, chickpea seedlings were placed on top of a metal to ensure that only the root was in direct contact with the liquid medium. Seedlings were maintained in the magenta boxes under long-day conditions (16 h light, 8 h dark; 25°C; 70% humidity), under slow agitation (20 rpm) until the time of harvest. Lower 50% of the roots from the mock and ABA treatments were harvested at nine different timepoints: 15, 30 and 45 minutes, and 1 h, 2 h, 4 h, 6h, 8 h and 24 h after ABA exposure. For each of the four biological replicates three roots were sampled. Samples were collected in 2-ml tubes containing ceramic beads and ground using a tissuelyser (Retsch MM300). The mRNA extraction and library preparation were performed according to a non-strand specific random primer-primed RNA-seq library protocol (Townsley et al. 2015). The libraries were pair-end sequenced (150 bp) using NovaSeq X Plus.

### RNA-seq mapping, data analysis

Paired-end reads were quality trimmed using Trim Galore (version 0.6.6) (Krueger 2015) using default settings. After trimming, reads were pseudo-aligned to the CDC Frontier chickpea reference transcriptome (Phytozome, https://phytozome-next.jgi.doe.gov/) using Kallisto (version 0.46.2) (Bray et al. 2016) with the following parameters:-b 100. DEGs were identified using the R package limma according (Ritchie et al. 2015) to (Jo et al. 2025). For every time point, we compared the ABA and mock treated samples using edgeR (Robinson et al. 2010) and DEGs were determined as genes with an Benjamini-Hochberg adjusted P value < 0.05 and at least 2-fold changes between treatments (Supplementary Table S2). DEGs were clustered based on their logFC values across all time points using kmeans function from R (k=8) (Supplementary Table S2). Annotation of DEGs was based on annotation info for the CDC Frontier genome on Phytozome.

### Promoter motif enrichment analysis

DNA motif enrichment analysis was performed as described (Jo et al. 2020), using the DNA motifs: GBOX (CACGTG), MYB41 (ACCTA), ATHB7 (ATNATTG), bHLH122 (CNACTTG), NF-YA (CCAAT), ANAC058 (CNTNNNNNNNANG), WRKY40 (TTGACT) and HAT5 (AATAATT) to screen the promoter regions (500 bps upstream the ATG) of DEGs identified in modules I, II, III, IV, V and VI. In summary, motifs were screened for enrichment using HOMER (homer.ucsd.edu/homer/motif/index.html) using the known function. For each DEG cluster, we generated 1000 sets of randomly selected genes that were of comparable numbers. Each set of promoters from each cluster were compared against the promoters of the respective 1,000 sets of random genes to calculate the P value significance of enrichment. P values were adjusted using the Bonferroni method, and a significance threshold of P < 0.01 was used to identify significantly enriched DNA motifs.

### Phylogenetic tree construction

Phylogenetic trees were generated using the methods described in (Jo et al. 2025). In summary, full length amino acid sequences of chickpea ABA-upregulated MYBs (Supplementary Fig. 4a), WRKYs (Supplementary Fig. 4b) or the best five BLAST hits of lignin-regulating MYBs (Supplementary Fig. S11f-i) from Phytozome v 1.0 and all annotated Arabidopsis MYB or WRKY TFs from TAIR10 were aligned using MAFFT (https://mafft.cbrc.jp/alignment/server/). The phylogenetic tree was determined using the maximum likelihood method with 1000 bootstraps using iQ-TREE 3.1 (http://iqtree.cibiv.univie.ac.at/) (Wong et al. 2025).

### Chickpea root protoplast isolation and scRNA-seq

For the scRNA-seq analysis we used protoplasts from roots (3 cm above the root tip, which corresponds to the middle-point of the root) of 3-day old chickpea seedlings grown in control conditions (mock) and in the presence of 2 µM ABA for 24 h. Approximately 15 roots were collected, cut into 1-2 mm slices and vacuum infiltrated for 15 minutes immersed in an enzyme solution consisting of 0.4 M D-Sorbitol, 20 mM KCl, 20 mM MES, 10 mM CaCl₂, 0.1% (w/v) BSA, 1.5% (w/v) Cellulase R10 (Duchefa), 0.5% (w/v) Macerozyme R10 (Duchefa), 5% (v/v) Viscozyme® L (Sigma) and 3.56 mM of 2-Mercaptoethanol. Roots were kept in the enzyme solution for 2 h under 40 rpm, and the resulting protoplast solution was filtered through a 40 µm mesh and centrifuged for 5 min at 200 x g at 4°C in a swinging bucket centrifuge with the de-acceleration set to minimal. Protoplasts were resuspended in 2 mL of resuspension solution (0.4 M D-Sorbitol, 20 mM KCl, 20 mM MES, 1 mM CaCl₂, 0.1% (w/v) BSA, 3.56 mM of 2-Mercaptoethanol). Chickpea root protoplasts were purified using a three-phase density gradient using Optiprep™ (Stemcell™ Technologies). An Optiprep™ working solution (WS) was prepared by mixing 0.06 g of KCl into 10 mL of Optiprep™. The density gradient was prepared in a 5 mL tube as follows: in the bottom layer, 2 mL of protoplast cell suspension was gently mixed and homogenized with 500 µL of OptiPrep™ WS (final density: 1.064 g/mL). For the middle layer, 1 mL of a mixture of 2 mL of resuspension buffer with 400 of OptiPrep™ WS (final density: 1.053 g/mL) was gently pipetted at the top of the bottom layer. For the topmost layer, 200 µL of the resuspension solution was carefully pipetted on top of the middle layer. The prepared gradient was centrifuged at 200 x g at 4°C for 5 minutes using a swinging bucket centrifuge. Following centrifugation, the protoplasts, concentrated in the topmost layer, were carefully collected, centrifuged again for 5 min at 200 × g at 4°C and resuspended to a final concentration of ∼1,000 cells per μl in resuspension solution. The protoplast suspension was loaded into microfluidic chips (10X Genomics) and subsequently processed using the 10x Chromium Controller from 10x Genomics, Single-cell libraries were constructed using the Single Cell 3′ v3.1 Kit (10x Genomics) following manufacturer instructions. Sequencing was performed using NovaSeq X Plus (PE150).

### Single-cell transcriptome analysis

Reads were mapped using CellRanger (10X Genomics) to the CDC Frontier chickpea reference genome (Phytozome, https://phytozome-next.jgi.doe.gov/) with appended chloroplast genome of chickpea (NC_011163.1) and the mitochondrial genome of *M. truncatula* (NC_029641.1). For an improved alignment, a modified version of the current Carietinum_492_v1.0 (Phytozome) was used. Due to the 3′ bias of sequencing reads generated by the 10X Single Cell 3′ v3.1 library kit, the 3′ UTR regions of all genes in the chickpea genome were expanded by 300 bps using BEDTools (Quinlan and Hall 2010), as the Carietinum_492_v1.0 gene model lacks annotated 3′ UTR regions. Before analysis in Seurat (Hao et al. 2024), the fraction of background reads was removed using SoupX (Young and Behjati 2020), cell doublets were removed using scDblFinder, and low-quality cells were identified and removed based on the number of detected features (< 500 features) and mitochondrial and chloroplast gene counts (> 5% of mitochondrial and chloroplast counts), which resulted in the detection of 10330 and 8724 high-quality cells in the ABA-and mock-treated root cells, respectively. Seurat objects from the ABA and mock samples were integrated using the IntegrateLayers () function from Seurat using the canonical correlation analysis (CCAIntegration) method (Butler et al. 2018). UMAP were calculated using 18 principal component dimensions and cell-clusters were identified using the Seurat ‘FindClusters’ function with the resolution parameter set to 0.7. Markers for each UMAP cluster were identified using the using the Seurat ‘FindAllMarkers’ and filtered for an average log2FC greater or equal than 0.5, adjusted P values lower or equal 0.01, percentage of cells within the cluster that expresses the gene greater or equal 20% and percentage difference to other clusters greater than 20%. The identities of specific UMAP clusters were annotated by comparing whether orthologues of marker genes were characterized as marker genes in the Arabidopsis scRNA-seq root database (scPlantDB) (He et al. 2024) and those empirically determined from the Arabidopsis scRNA-seq root atlas (Shahan et al. 2022). Additionally, the exodermis and endodermis cell clusters were identified based on the expression of a manually curated list of suberin biosynthesis genes and Casparian strip genes (Supplementary Table S3), respectively.

For the DEG analysis in each single-cell cluster, we compared the expression of genes between cells of the ABA treated roots and the mock treated roots. We use the ‘FindMarkers’ function from Seurat with the method MAST (Finak et al. 2015). We consider DEGs every gene with a log2FC equal or greater than 0.25 and an adjusted P value equal or lower of 0.01. The full list of cluster DEGs can be found in Supplementary Table S3.

### Lineage detection and pseudotime trajectory analysis

Lineage detection and pseudotime trajectory analysis were performed using the cell clusters representing the outer ground tissue (Outer Cortex 1 & 2 and Exodermis). The Meristem_1 cluster was included to anchor the cell lineage detection and pseudotime estimations. Lineage detection and pseudotime trajectory were detected using the Slingshot R package (Street et al. 2018) with the ‘getLineages’ and ‘slingshot’ functions, respectively.

### HCR Gold RNA-FISH

*C. arietinum* ‘Annigeri-1’ seeds were sterilized and germinated as described above. Following 3 days of germination in darkness, seedlings were transferred to light conditions for an additional 24 h (16 h light/8 h dark; 25°C; 70% relative humidity) in ½ MS liquid medium supplemented with 2 µM ABA. Following ABA treatment, roots were cut transversely at their midpoint, and 2-cm segments immediately above the cut were collected. Each segment was subsequently bisected longitudinally using a scalpel and immediately submerged in FAA fixative (3.7% formaldehyde, 5% glacial acetic acid, and 50% ethanol in RNase-free water) in a 12-well plate containing strainers. Samples were vacuum-infiltrated five times for 5 min each, until the root segments had completely sunk, and subsequently incubated for 1 h at room temperature, followed by overnight incubation at 4°C. Fixed root segments were dehydrated through successive 10-min incubations at room temperature in 70%, 90%, and twice in 100% ethanol. Samples were then washed twice in 100% methanol for 10 min each, incubated in methanol for an additional 1 h, and washed twice in 100% ethanol for 10 min each. Clearing and permeabilization were performed by incubating the samples in a 1:1 (v/v) mixture of Histo-Clear and ethanol for 30 min, followed by two 10-min washes in ethanol and three 5-min washes in methanol. Samples were subsequently rehydrated through a graded methanol series in DPBS-T (0.1% [v/v] Tween-20 in 1× DPBS), consisting of 75%, 50%, and 25% methanol, followed by two washes in methanol-free DPBS-T (0% methanol). All rehydration steps were performed for 5 min at room temperature. Partial cell wall digestion was performed by incubating the samples for 30 min in 1× cell wall digestion solution prepared in DPBS-T from a 6× stock containing 5 mg/mL macerozyme R-10 (Duchefa Biochemie, M8002), 5 mg/mL cellulase R-10 (Duchefa Biochemie, C8001), 2.5 mg/mL pectolyase Y-23 (Duchefa Biochemie, P8004), and 7.5 mg/mL pectinase (Sigma-Aldrich, 89216-50G). Samples were subsequently washed three times for 2 min each in DPBS-T and post-fixed for 45 min in fixative solution containing 3.7% formaldehyde (prepared by diluting a 37% formaldehyde stock solution to 10% [v/v] in DPBS-T). Following two 5-min washes in DPBS-T, samples were incubated with Proteinase K (80 µg/mL final concentration) in Proteinase K buffer (0.1 M Tris-HCl, pH 8.0, and 0.05 M EDTA) for 30 min at 37°C. After two additional 5-min washes in DPBS-T, samples were fixed again in fixative solution for 30 min at room temperature.

HCR Gold RNA-FISH probes and reagents were obtained from Molecular Instruments, and assays were performed with minor modifications to the manufacturer’s protocol. Pre-hybridization was performed by incubating the samples in probe hybridization buffer under vacuum (−0.04 MPa) for 10 min, followed by incubation at 37°C for 50 min. The probe solution was prepared by adding 20 µL of probe mixture to 1 mL of pre-warmed probe hybridization buffer (37°C). The pre-hybridization buffer was replaced with the probe solution, and samples were incubated overnight (∼20 h) at 37°C in a humidified chamber to prevent evaporation. Excess probes were removed by four 15-min washes in pre-warmed 1× probe wash buffer at 37°C, followed by two 10-min washes in 5× SSC-T (25% [v/v] 20× SSC and 0.1% [v/v] Tween-20) at room temperature. Samples were then incubated in amplification buffer under vacuum (−0.04 MPa) for 10 min, followed by incubation for 50 min at room temperature. Meanwhile, fluorescently labeled hairpin amplifiers (X1 and X2) were snap-cooled separately in PCR tubes by heating at 95°C for 90 s and subsequently cooling to room temperature in the dark for 30 min. The amplifier solution was prepared by adding 20 µL of each snap-cooled hairpin amplifier to 1 mL of amplification buffer. The pre-amplification buffer was replaced with the amplifier solution, and samples were incubated overnight (∼20 h) at room temperature in the dark. Following amplification, samples were washed four times for 15 min each in 1× amplifier wash buffer at room temperature in the dark, followed by two 20-min washes in 5× SSC-T at room temperature. Calcofluor White (0.5 mg/mL) was included in the second SSC-T wash to stain cell walls. Finally, samples were cleared in ClearSee for 3 days at RT and stored in the dark until imaging. For imaging, samples were mounted on microscope slides with the longitudinally cut surface facing upward and covered with coverslips. HCR probes targeting *Ca_20281* and *CaLTPG21* (*Ca_11846*) were designed for detection using the X1 and X3 amplifiers conjugated to Alexa Fluor 488 and Alexa Fluor 633, respectively. Images were acquired using a Zeiss LSM 880 confocal laser-scanning microscope equipped with a 20× air objective (Plan-Apochromat 20×/0.8 M27). Calcofluor White, Alexa Fluor 488, and Alexa Fluor 633 were excited using a 405-nm diode laser, a 488-nm Argon laser, and a 633-nm HeNe laser, respectively. For *Ca_20281*, fluorescence emission was collected at 411–474 nm for Calcofluor White and 508-552 nm for Alexa Fluor 488. For *CaLTPG21* (*Ca_11846*), fluorescence emission was collected at 410–523 nm for Calcofluor White and 638–747 nm for Alexa Fluor 633. The two fluorescence channels were acquired sequentially for each probe.

### Generation of hairy root transgenics and promoter fusions

Promoter regions (2 kb) were amplified from genomic DNA of chickpea cultivar Annigeri-1 using promoter-specific primers (Supplementary Table S4) and cloned into pENTR/D-TOPO vector (Invitrogen, USA). Positives clones were subsequently transferred into the Gateway-compatible destination vector pKGW-RedRoot, upstream of the *DsRed* reporter gene, using LR Clonase II (Invitrogen, USA). The resulting promoter-reporter constructs (*Ca11228p::DsRed* and *Ca00232p*::DsRed) were transformed into *Agrobacterium rhizogenes* strain ARqua1. Composite transgenic hairy roots were generated in chickpea cultivar BG 256 following previous methods with minor modifications (Mandal et al. 2020; Singh et al. 2020). ARqua1 strains carrying the constructs were first cultured in 5 mL LB with appropriate antibiotics at 28°C and 200 rpm for 24 hours. The 0.1% of primary culture was inoculated in 100 mL LB and grown until OD reached 0.8-1.0. Bacterial cells were pelleted at 5,000 x g for 10 minutes, resuspended in 500 μL LB and plated on LB agar. The plates were incubated at 28°C for 16-24 hours to produce a dense bacterial lawn. Roots of three-day-old chickpea seedlings were cut at a slant at the hypocotyl-root junction using a sterile scalpel, and the cut surfaces were rubbed on the bacterial lawn. Inoculated seedlings were transferred to Fähraeus medium (Singh et al. 2020), with the upper half of the media removed for shoot growth. The lower portions of the plates were covered with aluminum foil and incubated at 22°C for 3-4 weeks. Transformed roots were identified using a fluorescence microscope (Leica) and positive roots were longitudinally sectioned at 200 µm using a vibratome and imaged on a Zeiss LSM880 Airyscan confocal microscope with excitation at 561 nm and detected at range 570-640 nm.

### Constructs for transactivation assays: promoters

Promoter regions (∼2 kb upstream of the start codon) were PCR-amplified from *C. arietinum* Annigeri-1 genomic DNA using the primers listed in Supplementary Table S4. The amplified fragments were first cloned into the Gateway entry vector pDONR207 and then recombined into the Gateway-compatible destination vector pGreenII 0800-LUC. In the case of *pCa_01018*, the sequence was obtained pre-cloned in the pDONRZeo vector (GeneArt, Thermo Fisher Scientific) and was recombined directly into the destination vector pGreenII 0800-LUC. The sequence used is listed in Supplementary Table S4.

### Constructs for transactivation assays: transcription factors

Total RNA was extracted from roots of *C. arietinum* Annigeri-1 grown in ½ MS agar plates in the presence of ABA (2 μM) using the RNeasy Mini Kit (Qiagen), followed by DNase I treatment (Thermo Fisher Scientific). cDNA synthesis was performed using the RevertAid RT Reverse Transcription Kit (Thermo Fisher Scientific) with random hexamer primers. Coding sequences of the TFs were amplified using primers listed in Supplementary Table S4, cloned into the Gateway entry vector pDONR221, and subsequently recombined into the destination vector pH7WG2.

### Transient expression assays in N. benthamiana agroinfiltrated leaves

Competent *Agrobacterium tumefaciens* AGL-1 cells were transformed with the expression constructs. Single colonies were cultured in 20 mL LB medium at 28°C for 2 days with shaking. After reaching the desired OD600, the cultures were pelleted and resuspended to a final OD600 of 0.5 in half-strength MS medium (Duchefa Biochemie) supplemented with 10 mM MES hydrate (Sigma-Aldrich), 20 g/L sucrose (Sigma-Aldrich), and 200 μM acetosyringone (Sigma-Aldrich), adjusted to pH 5.6. The suspensions were incubated in darkness for 3–4 hours.

Agroinfiltration was performed on the abaxial side of 4-week-old *N. benthamiana* leaves using a 1 mL syringe. *Agrobacterium* strains carrying the promoter and transcription factor constructs were mixed prior to infiltration. Plants were maintained under normal light conditions. One day post-infiltration, the abaxial and adaxial sides of the infiltrated leaves were brushed with a solution containing 0.01% (v/v) Tween-20 and either 20 mM ABA (in 70% ethanol) diluted in water (final concentration of 75 µM) or a mock solution (equal volume of 70% ethanol in water). After overnight incubation (16–18 h), leaf sections from the infiltrated regions were collected for transactivation analysis. Firefly and Renilla luciferase activities were determined using the Dual-Luciferase Reporter Assay System kit (Promega) and measured using the GloMax Navigator microplate luminometer reader (Promega). Each transactivation assay was performed with 3 or 4 biological replicates at two independent experiments (Fig. 7b and Supplementary Fig. S17).

## Results

### Galegoid legumes have diverse root barrier phenotypes

To investigate the root suberization and lignification in galegoid legumes and the effect of ABA on it, we grew eight species selected based on previous reports of exodermis in legumes (Perumalla et al. 1990b; Schreiber et al. 1999; Taleisnik 1999; Bani et al. 2018; Sharma et al. 2023) in the presence or absence of 2 mM ABA (Fig. 1b). After 3 days, root cross-sections of the middle region of the root (50% region) were stained with Fluorol Yellow (FY) and Basic Fuchsin (BF), fluorescent dyes that bind to suberin and lignin, respectively (Heslop-Harrison et al. 1986; Brundrett et al. 1991; Dharmawardhana et al. 1992; Lux et al. 2005), and the fluorescence intensity of the cell wall was quantified across individual cell layers (Fig. 1c, Supplementary Fig. S1). Based on this histochemical screen, ABA induced suberization and lignification in different root cell layers across the galegoid clade (Fig. 1c, Supplementary Fig. S1-S2). In chickpea and *L. oleraceus*, ABA promoted accumulation of suberin and non-polar lignin specifically in the outermost cortical cell layer (C1) and not in the endodermis (EN), indicating presence of an exodermis in these species. In contrast, the rest of the species did not have a strong indication of exodermis. For these, ABA induced the suberization of the endodermis in *Lens culinaris* (lentil), *Medicago truncatula* (barrel medic), *Medicago sativa* (alfalfa), and *Trifolium repens* (white clover), with a weak FY and BF signal observed in cortical cell layers. In *Lotus japonicus* (narrowleaf trefoil), the process of root suberization in the endodermis occurred independently of exogenous ABA at the sampled developmental stage. We indeed observed that the root zone (developmental age) affected the number of suberized cells (Supplementary Fig. S3) and that the ABA treatment slowed down root growth (Supplementary Fig. S4) across species. *Vicia sativa* (common vetch) had unique FY and BF patterns in both outer cortex and endodermis: multiple outer cortex layers responded to ABA, and strong non-polar FY and BF signal was detected in the endodermis in both treatments. All other species exhibited anticlinally localized BF signal in endodermis in both mock and ABA treatments (Supplementary Fig. S2), in line with a lignified Casparian strip under developmental control.

### Chickpea forms a functional exodermal barrier in response to ABA

Suberin monomer profiling with gas chromatography-mass spectrometry (GC-MS) further confirmed that ABA promotes the accumulation of suberin monomers in chickpea, *M. truncatula*, and *V. sativa* roots (Fig. 2a, b, Supplementary Fig. S5a, Supplementary Table S1), with the overall increase of *cis-* and *trans*-ferulic acid, long-chain fatty acids (C18, C20, C22), α-ω-diacids (C16, C18:1, C18) and ω-hydroxy acids (C16, C18:1 and C22) (Supplementary Fig. S5a). As the cortex layers are not separated from endodermis for these analyses, for *M. truncatula* and *V. sativa*, these include the increases in endodermal suberin. Given these diverse patterns of suberin accumulation, we then examined the ultrastructure of the suberin in C1 layer (Fig. 2c) and endodermis (Supplementary Fig. S5b) using transmission electron microscopy (TEM) for the same three species with contrasting FY phenotypes. In chickpea roots grown in the presence of ABA, C1 displayed a clear SL, characterized by a pattern of alternating dark and light bands (Falk and El-Hadidi 1961) (Fig. 2c). In contrast, while the SL was clearly present in the endodermal cell layer of *M. truncatula* and *V. sativa* (Supplementary Fig. S5b), it was absent in the C1 cells in these species (Fig. 2c). Despite the high FY signal detected in the outer cortex of *V. sativa* (Fig. 1c), TEM revealed an amorphous suberin structure lacking the alternating banding pattern typical for SL (Fig. 2c).

**Figure 2.**
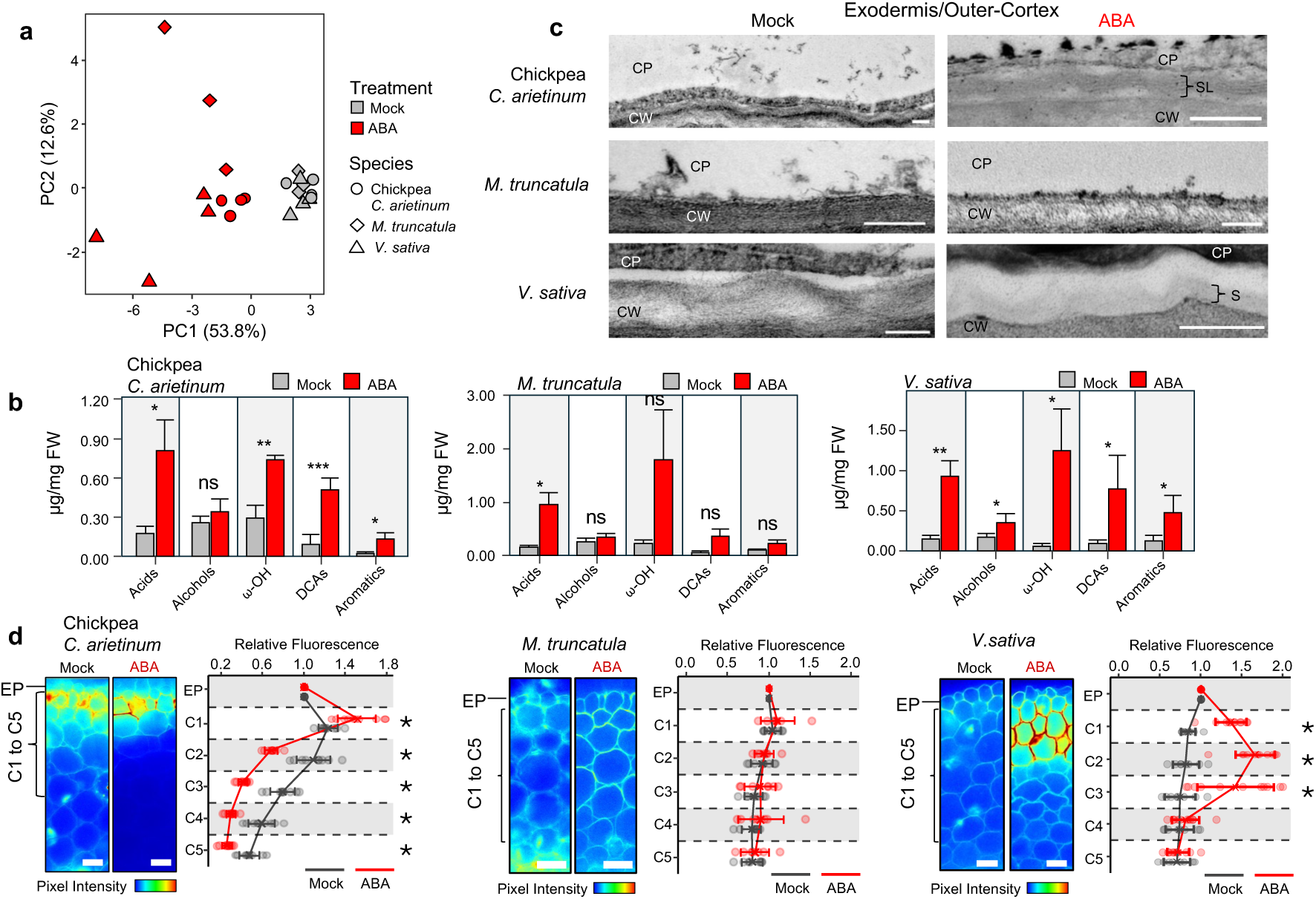
Biochemical, structural and functional evaluation of ABA-induced cell wall modifications show that chickpea forms an exodermis with a suberin lamella and an apoplastic diffusion barrier. **a,** Principal Component Analysis (PCA) of suberin monomer abundances in roots of chickpea*, M. truncatula* and *V. sativa* treated with mock or 2 µM ABA for 24 hours. **b,** Abundance of major classes of suberin monomers in roots treated with mock (grey) or 2 µM ABA (red) for 24 hours. Asterisks indicate significance based on Student’s t-test with a P-value threshold of 0.05 (n = 4). **c,** Outermost cortex cell wall shown in transmission electron microscopy (TEM) images of root cross-sections at the 75% region of Chickpea, *M. truncatula* and *V. sativa* roots treated with mock or 2 µM ABA for 3 days. The cell wall (CW) depicted in the images refers to the cell wall between two adjacent cells of the exodermis (in chickpea) or in the outermost cortex layer (in *M. truncatula* and *V. sativa*). Scale bar, 200 nm. SL: suberin lamellae; S: suberin; CW: cell wall; CP: cytoplasm. **d,** Propidium Iodide (PI) diffusion assay of chickpea, *M. truncatula* and *V. sativa* roots after treatment with mock (black) or 2 µM ABA (red) for 3 days. For each species, the left image shows a representative image of laser scanning confocal microscopy for PI in root cross-sections, with the PI signal indicating diffusion across epidermal and cortex cells. Scale bar, 25 µm. On the right, line plots show the relative pixel intensity of the PI dye cell wall in different layers of the root cortex (C1 to C5) relative to the intensity of the signal detected in the epidermis (EP). Asterisks indicate significance based on Wilcoxon rank-sum test with statistical significance of P < 0.05 (n ³ 6).

To determine if ABA-induced compounds in the cortex of these species function as an apoplastic diffusion barrier, we tested if they restrict the diffusion of propidium iodide (PI) (Alassimone et al. 2010; Manzano et al. 2024). PI apoplastic diffusion was blocked in chickpea roots treated with ABA as indicated by the lack of PI fluorescent signal in inner cortical cell layers when compared to mock-treated roots (Fig. 2d). This suggests that ABA-induced cell-type-specific modifications in the cell wall of C1 results in the establishment of a functional apoplastic barrier in chickpea. This supports the chickpea C1 being a functional exodermis. However, as chickpea exodermis has both SL and lignin in response to ABA, we cannot determine which one(s) provide the apoplastic barrier function. Previous work supports lignin, the main component of the Casparian strip, critical for blocking apoplastic diffusion of PI (Alassimone et al. 2010; Naseer et al. 2012).

In *M. truncatula* and *V. sativa*, PI diffused into the inner cortex layers in both mock and ABA-treated roots (Fig. 2d). The cortex layers with FY and BF signal also resulted in strong PI signal in chickpea and *V. sativa* (Fig. 1c, Fig. 2d), which may relate to cross-reactiveness of PI with the cell wall modifications. In summary, our results demonstrate that the stress hormone ABA promotes suberin and lignin accumulation in the outer cortex of various legume species. Specifically, chickpea exhibited a robust single-layer response with deposition of structured SL and lignin and forming a functional apoplastic diffusion barrier, i.e., an exodermis.

### ABA-induced transcriptome changes specify temporally and functionally distinct expression modules in chickpea roots

The suberin biosynthesis and deposition pathway is a coordinated process with many enzymes that operate in different cellular compartments (Serra and Geldner 2022) (Fig. 3a). To understand how the process of ABA-induced exodermal differentiation leading to apoplastic barrier formation is transcriptionally regulated, we profiled the chickpea root transcriptome at nine different timepoints: 15, 30 and 45 minutes, and 1h, 2h, 4h, 6h, 8h and 24h after mock or ABA exposure (Fig. 3b, Supplementary Table S2). ABA induced primarily upregulation of gene expression (Fig. 3c). K-means clustering of differentially expressed genes (DEGs) identified temporal expression patterns in response to ABA (Fig. 3d, Supplementary Fig. S6a, Supplementary Table S2). Genes exhibiting a rapid upregulation (30 minutes to 1 hour) were grouped into Modules I, II and III (“early”), while Modules IV, V and VI represented temporally delayed transcriptional responses showing continuous upregulation of genes from 2h or later (“late”, Fig. 3d, Supplementary Fig. S6a). Genes involved with the canonical ABA response, like members of Clade A *PROTEIN PHOSPHATASE TYPE 2C* (*PP2C*), *RESPONSIVE TO DESICCATION 29* (*RD29*), and *LATE EMBRYOGENESIS ABUNDANT* (*LEA*) were found in the early modules (Fig. 3d, e, Supplementary Fig. S6b). In contrast, suberin biosynthesis and deposition genes (Fig. 3a), were found in the late modules (Fig. 3c, f, Supplementary Fig. S6c). Matching the transcriptional timing, suberin accumulation in the exodermis could be detected by FY after 24h (Supplementary Fig. S7). Contrastingly, lignin biosynthesis and deposition genes were found in both early (II, III) and late (V, VI) clusters (Supplementary Fig. S6d, e). In summary, the expression of suberin biosynthesis genes is temporally separated from genes in the general ABA response in chickpea roots, while lignin overlaps with both, indicating separate regulatory mechanisms.

**Figure 3.**
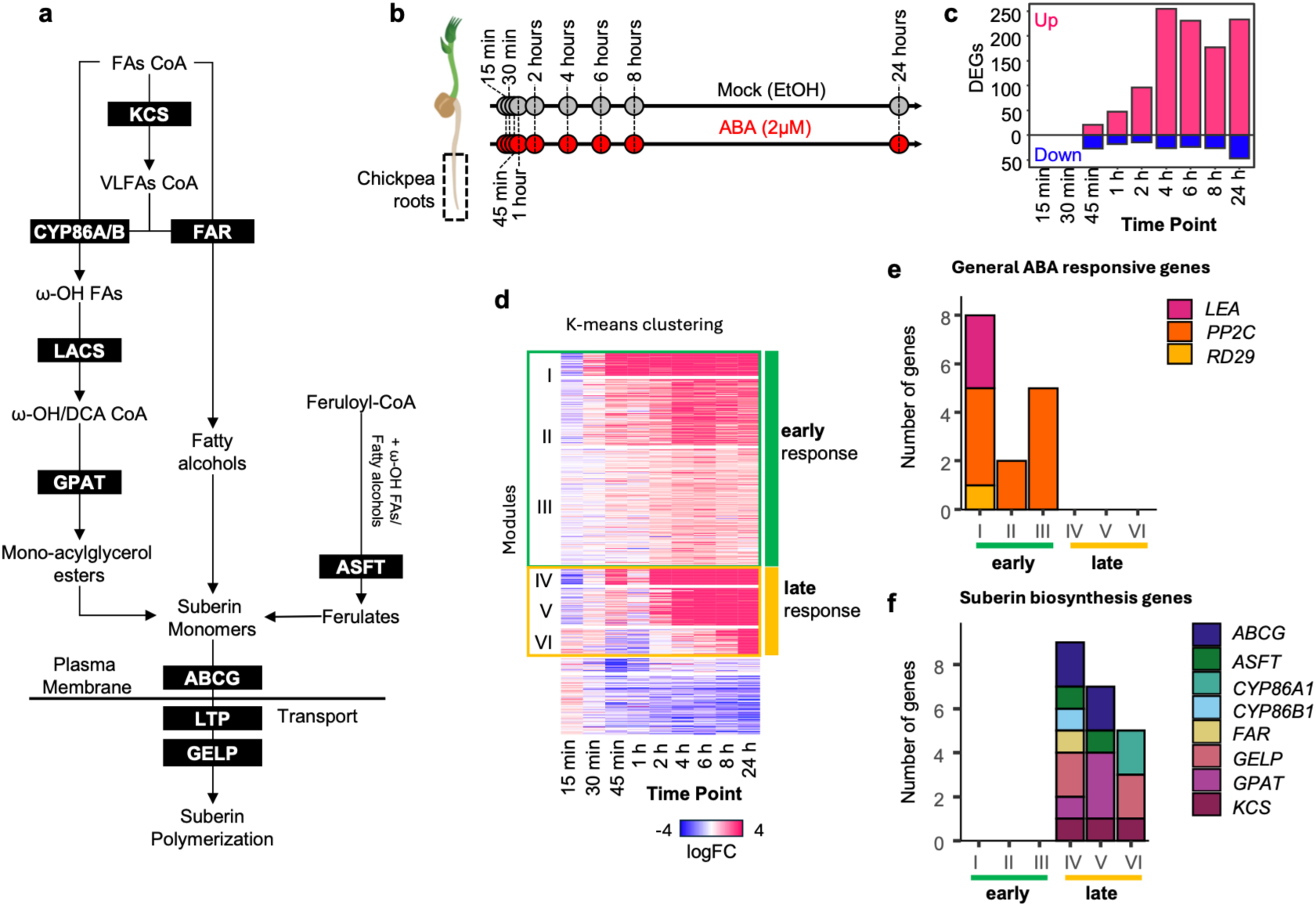
ABA induces temporally separated general ABA response and suberin biosynthesis programs in chickpea roots. **a,** Simplified schematic diagram of suberin biosynthesis, transport and polymerization adapted from (Cantó-Pastor et al. 2024). Black boxes indicate enzymes or transport proteins involved in each step of the pathway. FAs: fatty-acids; VLCFA: very-long fatty-acids; KCS: 3-ketoacyl-CoA synthase; FAR: Fatty acyl-CoA reductase; CYP86A/B: cytochrome P450, family 86, subfamily A or B; LACS: long-chain acyl-coenzyme A synthetase; GPAT: glycerol-3-phosphate acyltransferase; ASFT: Aliphatic suberin feruloyl-transferase; ABCG: ABC transporter G; LTP: lipid transfer protein; GELP: GDSL esterase/lipases. **b,** Schematic representations of time points (15, 30 and 45 minutes (min) and 1, 2, 4, 6, 8 and 24 hours (h) used to profile the transcriptome changes of roots (lower half of the root) from 3-day old chickpea treated with mock (grey) and 2 µM ABA (red). **c**, Number of differentially expressed genes (DEGs) at each time point for ABA-treated roots compared with mock. DEGs were determined as genes with an Benjamini-Hochberg adjusted P value ≤ 0.05 and log2FC ≥ 1 (upregulated, pink) or log2FC ≤-1 (downregulated, blue). The full lists of DEGs are given in Supplementary Table S2. **d**, Temporal modules detection using K-means clustering of DEGs (k = 8). The heatmap shows logFC values of each gene at each timepoint. For further analysis, we focused on six modules that had the highest upregulation in response to ABA (Modules I, II, III, IV, V and VI). These modules were grouped into two groups: early (I, II and III) and late (IV, V and VI) response to ABA based on their temporal profile (Supplementary Fig. S7a). The list of genes in temporal modules is given in Supplementary Table S2. **e**, Number of chickpea genes related to the canonical ABA response identified in modules I-VI. *LEA*: *LATE EMBRYOGENESIS ABUNDANT*; *PP2C*: *PROTEIN PHOSPHATASE 2C*; *RDB29B*: *RESPONSIVE TO DESICCATION 29B*. **g**, Number of chickpea genes related to suberin biosynthesis, transport and polymerization (listed in **a**) in modules I-VI.

To understand the transcriptional regulatory mechanisms underlying the temporal separation between the general ABA response and the suberin biosynthesis program, we analysed the DNA *cis*-element composition of promoters (500 base pairs upstream the coding sequence) of genes belonging to modules I-VI (Fig. 4a, b, c, Supplementary Fig. S8). Motif enrichment analysis revealed that the G-box *cis*-element (CACGTG), the main DNA element targeted by the canonical ABA response (Guiltinan et al. 1990; Choi et al. 2000) was significantly enriched only in the promoters of the early modules (Fig. 4a). This supports the rapid transcriptional response observed in chickpea roots occurring through the canonical ABA signalling pathway (Ng et al. 2014). We observed other *cis*-elements enriched in the promoters of late modules, including that of suberin-regulating MYB41 TF (ACCTA) (O’Malley et al. 2016; Zenker et al. 2024; Baumgart et al. 2025) (Fig. 4b) and WRKY40 (TTGACT) (O’Malley et al. 2016) (Fig. 4c). This indicates that albeit ABA-induced, suberin biosynthesis in the chickpea exodermis is not directly controlled by the canonical ABA signalling pathway that utilizes the G-box element. Instead, it appears to be regulated through ABA-induced changes in the regulatory state of exodermal cells, with the involvement of other TFs, like MYBs and WRKYs, that control the temporally separated expression of suberin biosynthesis genes.

**Figure 4.**
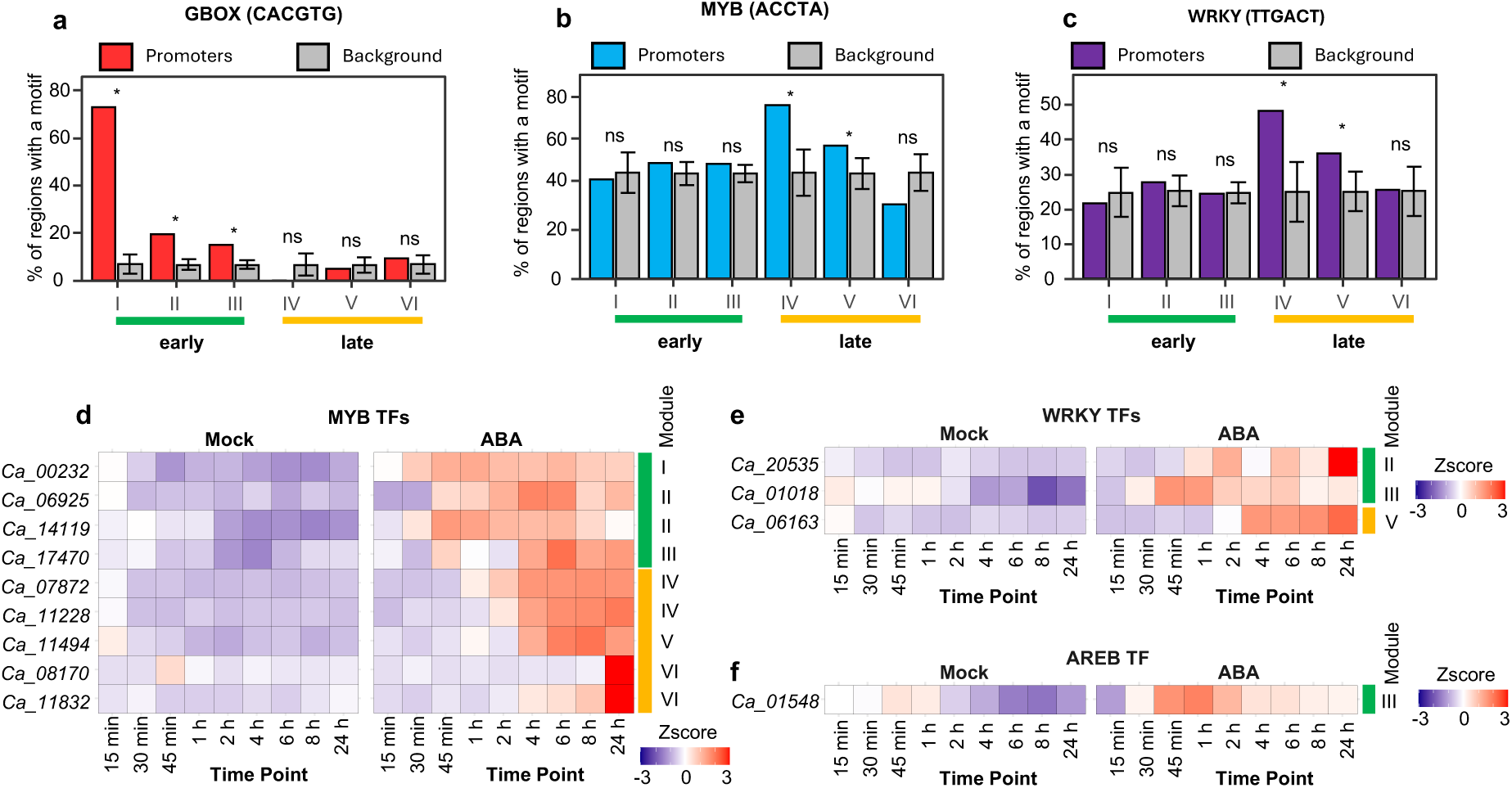
Putative mechanisms of the temporal distinction between ABA response and suberin biosynthesis program in chickpea roots. **a,** GBOX (CACGTG), **b,** MYB41 (ACCTA) and **c**, WRKY40 (TTGACT) motifs enrichment in the module I-VI promoters (500 base pairs (bps) upstream the start codon (ATG) of each gene). Bar graphs show the percentage of promoters of DEGs (colored) and 1000 sets of promoters of randomly selected genes with equivalent number to the compared DEG list (grey) containing the indicated DNA motifs. A Bonferroni-adjusted P-value threshold of 0.01 was used to indicate the statistical significance (*) of enrichment, while ns indicates no significance. Expression of **d,** *MYB*, **e**, *WRKY* and **f**, *AREB* DEGs. Heatmaps show relative mRNA levels of chickpea treated with mock and 2 µM ABA for 15, 30 and 45 minutes (min) and 1, 2, 4, 6, 8 and 24 hours (h). The temporal module corresponding to each gene is specified.

Based on these results, we hypothesized that ABA enhances the expression and/or activity of specific MYB and WRKY TFs through the canonical ABA signalling pathway. These TFs, in turn, initiate the suberin biosynthesis program in chickpea roots. Among the ABA-induced *MYBs* and *WRKYs*, *Ca_00232, Ca_14119* (Fig. 4d), and *Ca_01018* (Fig. 4e) were rapidly upregulated within 45 minutes of ABA exposure, a pace similar to *Ca_01548*, an *AREB* (*ABA RESPONSIVE ELEMENTS-BINDING PROTEIN*) and an indicator of the canonical ABA response (Fig. 4f). Phylogenetic analysis revealed that Ca_00232 shares a close evolutionary relationship with AtMYB74 and AtMYB102 and Ca_14119 with AtMYB41 (Supplementary Fig. S9a). In Arabidopsis, AtMYB74 has been characterized as direct target of the ABA signalling, and AtMYB41 regulates the suberization of the root endodermis (Kosma et al. 2014; Barberon et al. 2016; Shukla et al. 2021; Ortiz-García et al. 2022; Xu et al. 2022). For the WRKY Ca_01018, the closest homologs were AtWRKY23 and AtWRKY68 (Supplementary Fig. S9b). Other *MYB* and *WRKY* DEGs presented a later response to ABA in a similar temporal pattern as the expression of suberin biosynthesis genes (Fig. 4d, e, Supplementary Fig. S6c), and some of these, e.g. *Ca_11228*, *Ca_11494* and *Ca_11832*, are close orthologues of known regulators of suberin, such as Arabidopsis *SUBERMAN* (*AtMYB39*) (Cohen et al. 2020) (Supplementary Fig. S9a, Fig. 4d).

### Establishment of a single-cell atlas of the chickpea root in response to ABA

As suberin deposition in response to ABA is highly exodermis-specific in chickpea seedling roots, we next profiled the single cell transcriptome of chickpea roots in treated with either mock or 2 mM ABA for 24 hours to infer spatial expression patterns (Fig. 5a, Supplementary Fig. S10a, b). This timepoint allowed us to capture the differentiation trajectory until fully differentiated exodermis with initiation of the suberin and lignin biosynthesis program in the root exodermis. We profiled the transcriptome of over 8724 and 10330 good-quality cells (Supplementary Fig. S10c, d) isolated from mock and ABA treatments, respectively. After the integration of mock and ABA datasets (Fig. 5b), we detected 23 uniform manifold approximation and projection (UMAP) clusters representative of specific cell types of the chickpea root (Fig. 5c). To infer cell identities to each UMAP cluster, we compared the list of marker genes of each UMAP cluster to orthologues of marker genes characterized as marker genes in the Arabidopsis scRNA-seq root database scPlantDB (Fig. 5d) (He et al. 2024). In our dataset we considered marker genes to have expression in at least 20% of cells of the UMAP cluster, a percentage difference of at least 20% to other clusters, an average logFC greater than 0.5, and an adjusted P-value lower than 0.01 (Supplementary Table S3). We further analysed the expression of empirically determined root cell markers in Arabidopsis (Shahan et al. 2022) to refine the annotation of individual clusters, including xylem, cortex, phloem and epidermis (Supplementary Fig. S10e, f). For instance, two Xylem cluster marker genes, *Ca_17365* and *Ca_00412*, are orthologues of Arabidopsis *CELLULOSE SYNTHASE5* (*CESA5*) and *CESA8*, which were previously characterized as protoxylem markers (Shahan et al. 2022) (Supplementary Table S3). Similarly, Trichoblast cluster marker genes, such as *Ca_02397* and *Ca_18366*, correspond to Arabidopsis *EXPANSIN7* (*EXP7*) and *ROOT HAIR DEFECTIVE6-LIKE2* (*RSL2*), both well-established trichoblast marker genes (Shahan et al. 2022) (Supplementary Fig. S10f, Supplementary Table S3). Similarly, *Ca_23533,* an orthologue of the metaphloem and companion cell marker gene *WOODY* (*WDY*) (Shahan et al. 2022), was a marker gene of Phloem clusters 1 and 2 (Supplementary Fig. S10f, Supplementary Table S3). Marker genes of the meristem cell clusters (Meristem 1-4) were orthologues of genes in the meristem and dividing cells categories (G2-M phase, S phase, Meristematic cell) in the scPlantDB (Supplementary Table S3). For the annotation of the two barrier cell types, we leveraged the Casparian strip and its specialized deposition machinery as a unique cellular feature of endodermis (Supplementary Fig. S11a, b). Indeed, the orthologs of the genes involved in Casparian strip formation were exclusively expressed in the Endodermis cluster (Supplementary Fig. S11c), including *GASSHO1/SCHENGEN3* (*SGN3*), *CASPARIAN STRIP MEMBRANE PROTEINs* (*CASPs*) and their regulator *MYB36* (Roppolo et al. 2011; Kamiya et al. 2015; Liberman et al. 2015). Additionally, the Casparian strip formation in Arabidopsis requires the stele-produced peptide CASPARIAN STRIP INTEGRITY FACTOR1 (CIF1) (Doblas et al. 2017; Nakayama et al. 2017). We observed that also in chickpea, the closest orthologue to *AtCIF1* (*Ca_20263*) was highly expressed in stele cell clusters (Pericycle, Phloem 1&2, Stele Initials and Cambium) (Supplementary Fig. S11c). One of the UMAP clusters contained only three marker genes, it was designated as “unknown”. The full list of marker genes and their close orthologues in the Arabidopsis scPlantDB can be found in Supplementary Table S3. Overall, these annotated cell clusters provide an atlas of cell types of the chickpea root and offer valuable insights into the molecular mechanisms of root cell-type-specific responses to ABA.

**Figure 5.**
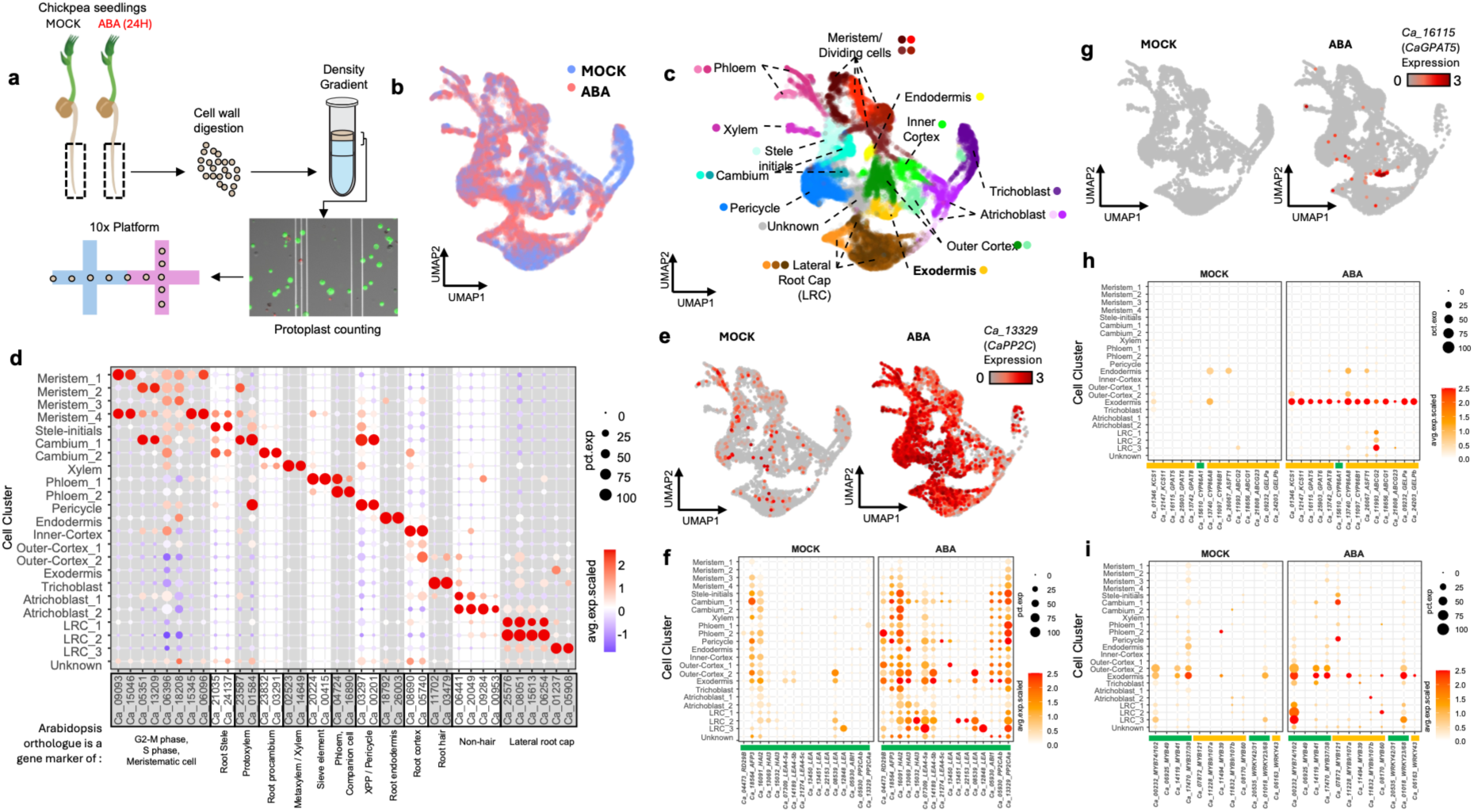
Single cell atlas of the chickpea root treated with ABA. **a,** Overview of the scRNA-seq experiment in chickpea roots. 3-day-old chickpea seedlings were transferred to ½ MS medium plates containing either 2 µM ABA or a mock treatment. After 24 hours, protoplasts were released from roots (3 cm root tip) and loaded into the 10X Genomics Chromium chip for making the single-cell transcriptome libraries. **b,** UMAP of integrated single-cell datasets from mock (lavender) and 2 µM ABA for 24h (salmon) treated roots. **c,** UMAP showing the twenty-three single-cell clusters of the chickpea root colored by their respective annotated cell identities. The list of marker genes for each single-cell cluster is presented in Supplementary Table S3. **d**, Expression of the top 2 marker genes for each cluster. Dot size indicates the percentage of cells in the cluster that expresses the specific gene (% expressed). Dot colors represent the average scaled expression of genes in the clusters (Avg. expression). The top 2 markers were identified by looking at the genes with the lowest adjusted P-values and the greatest percentage differences with other clusters. In the bottom of the dot plot, genes are annotated to indicate if the marker gene has an orthologue annotated as a marker gene in the described cell type in the Arabidopsis root scPlantdb (He et al. 2024). To filter for good quality data in the scPlantDB, we manually curated the datasets by only keeping datasets in which *AtMYB36* was annotated as a marker for the endodermal cell cluster. **e**, Feature plots of mock and ABA-treated cells showing the wide expression level of the ABA-responsive gene *Ca_13329* (*CaPP2C*) across multiple cell-types. **f**, Dot plot showing the expression of early ABA responsive genes in the single-cell data. **g**, Feature plots of mock and ABA treated cells showing the localized expression level of the suberin biosynthesis related gene *Ca_16115* (*CaGPAT5*) in the exodermis cluster. **h**, Dot plot showing the localized expression of suberin biosynthesis genes in the single-cell data. **i**, Dot plot of *MYB* and *WRKY* TFs identified as upregulated DEGs in the time course bulk RNA-seq experiment. For dot plots, the color next to the gene name indicates their early (green) or late (yellow) temporal module in the bulk RNA-seq dataset.

### Identifying the chickpea root cell-type specific responses to ABA

Across all cell types, we observed a slight overrepresentation of cells from ABA-treated roots compared to the mock control (Supplementary Fig. S10g, h), which likely reflects the higher total cells originated in the ABA-treated sample (Supplementary Fig. S10c). The only exceptions to this trend were the Trichoblast and Outer-Cortex_2 clusters, where most cells originated from the mock-treated roots (Supplementary Fig. S10g, h). The reduced cell representation in the Trichoblast cluster is consistent with the known role of ABA in repressing root hair differentiation (Rymen et al. 2017). Similarly, for the Outer-Cortex_2 cluster, this reduction may reflect a decreased trajectory of outer cortex cells differentiating into the exodermis. This indicates that the ABA treatment might selectively control the differentiation of specific cell types of the chickpea root.

When examining transcriptional changes in response to ABA in the single-cell dataset, we observed a response pattern consistent with that of the temporal bulk RNA-seq experiment (Supplementary Fig. S12a-d). This suggests that the overall transcriptional response in roots under the exposure of ABA is preserved in the individual cells of the chickpea root (Supplementary Fig. S11a-d). For instance, ABA early responsive genes, like the *Ca_13329* (*PP2C*) and *Ca_07309* (*LEA4-5a*), were shown to be expressed across multiple cell types of the ABA-treated sample, but not in the mock sample (Fig. 5e, f). This indicates that at the transcriptional level, most cell types in the chickpea root are capable of responding to ABA, suggesting that the early responses detected at the bulk level reflect a robust transcriptional response across major root cell types.

In accordance with what we observed at the cellular level (Fig. 1c) and transcriptional level (Fig. 3f), our single cell transcriptome data revealed the upregulation of suberin biosynthesis genes in response to ABA (Fig. 5g, h). More importantly, we observed that this occurred in a unique cell cluster of the chickpea root (Fig. 5g, h). We used this localized response as a proxy to identify the cell cluster that represents the root exodermis. Among the suberin biosynthesis genes, we identified one *CaASFT1* (*Ca_26067*), two *CYP86s* (*CaCYP86A1*: *Ca_15610*; *CaCYB86B1*: *Ca_11097*), two *KCSs* (*CaKCS1*: *Ca_01346*; *CaKCS11*: *Ca_12147*); one *ABCG* (*CaABCG23*: *Ca_21808*), four *GPATs* (*CaGPAT1*: *Ca_15259, CaGPAT5*: *Ca_16115, CaGPAT6*: *Ca_25903* and *CaGPAT8*: *Ca_13742*) and two *GELPs* (*CaGELPa*: *Ca_09232*; *CaGELPb*: *Ca_24203*) to be mainly expressed in the exodermis cluster in response to ABA (Fig. 5h). A similar cell-type transcriptional response was observed in another independent experiment (Supplementary Fig. S13a-g). Additionally, we experimentally validated this cluster as representing the exodermis. We selected transcripts enriched in clusters representing exodermis and its overlaying cell types and tested the accumulation of *Ca_20281* and *CaLTPG21* (*Ca_11846*) via hybridization chain reaction fluorescence *in situ* hybridization (HCR-FISH, Supplementary Fig. S14a-d) and *CaMYB74/102* (*Ca_00232*) and *CaMYB9/107* (*Ca_11228*) via promoter fusions expressed in chickpea hairy roots (Supplementary Fig. S14e-h). Expression profiles identified via both approaches were in alignment with cluster annotation. This suggests that we successfully distinguished the exodermis cell-type transcriptional responses from nearby clusters.

Because enzymatic cell wall digestion is known to trigger stress-related responses in plants (Tenorio Berrío et al. 2025), we investigated whether protoplasting induces the expression of suberin biosynthesis genes. To address this, we performed bulk RNA sequencing on root protoplasts and compared them with undigested whole roots (Supplementary Fig. S15a, b). This analysis identified 1,457 upregulated and 665 downregulated DEGs in the digested roots (Supplementary Fig. S15c, Supplementary Table S5). Examination of their cell-type expression patterns revealed that these genes are distributed across multiple cell types (Supplementary Fig. S15d). Furthermore, only a small fraction of cell-type specific DEGs in the single-cell data (Supplementary Table S3) appeared to be induced by enzymatic digestion (Supplementary Fig. S15e, f), and more importantly, none of the identified ABA-induced suberin biosynthesis genes in the exodermis were found to be upregulated in the digested roots compared to the undigested roots (Supplementary Table S3 and S5). These findings confirm that our characterized suberin transcriptional response accurately captures cell-type-specific responses to ABA in chickpea roots, rather than artifacts of stress caused by the protoplast isolation.

Finally, we also mapped the expression of ABA-induced lignin biosynthesis genes in the single cell data and observed expression in multiple clusters, especially those representing the cell types that use lignin in their secondary cell walls: exodermis, endodermis and xylem (Supplementary Fig. S11c, d). The lignin biosynthesis genes belonging to early temporal modules were expressed in cell clusters representing endodermis and exodermis (Supplementary Fig. S11c). Interestingly, while this expression pattern seems to be independent of exogeneous ABA in the endodermis (Supplementary Fig. S11c, d), many of these lignin biosynthesis genes seems to be expressed only in response to ABA (Supplementary Fig. S11d). For instance, genes *Ca_17252* and *Ca_12445* orthologues of the Arabidopsis *LACCASE3* (*LAC3*) and *LAC5*, respectively, shown to be expressed in the exodermis cell layer in response to ABA. These observations indicate that while cell wall lignification in the endodermis and exodermis involves some of the same genes, their upstream regulation is distinct: the endodermis is controlled by the canonical SGN3/MYB36 regulatory module, whereas lignification of the exodermis is regulated by an ABA-inducible gene regulatory module. Lignin is also known to be regulated by MYB TFs (Xie et al. 2018), so we queried the closest homologs of known lignin-regulating MYBs (Supplementary Fig. S11e-i) in our data. While none of these homologs were induced by ABA, *Ca_22597*, a homolog of the positive regulator *AtMYB63* (Zhou et al. 2009), showed expression in select cells of the exodermis cluster (Supplementary Fig. S11e).

Altogether, we provide a cell-type-specific overview of ABA responses across distinct cell types in the chickpea root. Importantly, we successfully captured the ABA-induced transcriptional changes underlying the accumulation of suberin and lignin in the cell walls of the chickpea exodermis, consistent with our previous cellular, biochemical, and bulk transcriptomic analyses. To facilitate data exploration, we developed a Shiny app for browsing gene expression across single-cell and bulk transcriptome datasets of chickpea roots in response to ABA. To simplify the visualization of single-cell data, we also integrated a pseudobulk expression profile into a representative map of the chickpea root cross-section using the ggPlantmap R package (Jo and Kajala 2024). The app is publicly available to allow the research community to freely explore the datasets generated in this study (https://kaisakajala-scannigeri.share.connect.posit.cloud/).

### Temporal and developmental trajectories of *MYB* and *WRKY* TFs align

To detect the TF regulatory network responsible for the ABA-inducible process of exodermis suberization, we queried the spatial expression of *MYBs* and *WRKYs* identified in the time-course experiment (Fig. 4d, e). In accordance to the general trends for early and late modules, we observed that TF genes from the early clusters, including *Ca_00232 (CaMYB74/102), Ca_14119 (CaMYB41)*, *Ca_17479 (CaMYB37/38)* and *Ca_01018(CaWRKY23/68*) were expressed in the exodermis cluster, but also in clusters representing cell types overlaying exodermis (Lateral Root Cap, Outer-Cortex and Trichoblast clusters) (Fig. 5i), while TFs from the late temporal modules, such as *Ca*_*11228* (*CaMYB9/107*) and *Ca*_*06163* (*CaWRKY43),* were specific to the exodermis cluster only in the ABA-treated roots (Fig. 5i). Altogether, these data may indicate a hierarchical cascade of ABA-induced TFs, where the downstream TFs are restricted into one cell type, the exodermis, to induce suberin biosynthesis in cell-type specific manner in the chickpea root. Since we observed that many of the suberin biosynthesis genes to be solely expressed in the exodermis cluster, we propose that chickpea exodermis is uniquely specialized to translate the ABA-mediated changes in the regulatory landscape into the activation of the suberin biosynthesis program.

To reconstruct the spatial and temporal dynamics of ABA-induced suberization in the exodermis, we utilized the Slingshot package (Street et al. 2018) to infer developmental trajectories of outer cortex and exodermis cells using the single-cell data of the ABA treated roots (Fig. 6a). Unsupervised lineage detection revealed two main lineages starting from the Meristem_1 cluster, Lineage 1 to Exodermis_1 and Lineage 2 to OuterCortex_2, both via OuterCortex_1 (Fig. 6b, c, Supplementary Fig. S16a). Pseudotime estimation in Lineage 1 revealed that the expression of suberin biosynthesis gene expression initiated late in the exodermal developmental trajectory (Fig. 6c), while they were not expressed in Lineage 2 (Supplementary Fig. S16b-d). This suggests that our pseudotime trajectory of Lineage 1 represents the developmental trajectory of ABA response along the exodermis in chickpea. When looking at the expression of TFs along the Lineage 1 pseudotime trajectory, we observed that the majority of *MYBs* upregulated by ABA showed expression patterns similar to the suberin biosynthesis genes (Fig. 6d). However, similarly to the time-course data (Fig. 4d, e), *CaMYB74/102 (Ca_00232)* and *CaWRKY23/68 (Ca_01018)* showed early expression patterns along the exodermal lineage trajectory which coincides with the onset of the general ABA response in the exodermis (Fig. 6e, f). Thus, our single-cell analysis led us to hypothesise a role for CaMYB74/102 (Ca_00232) and CaWRKY23/68 (Ca_01018) in coordinating the ABA early perception and the delayed spatial control of suberin biosynthesis in the chickpea root exodermis.

**Figure 6.**
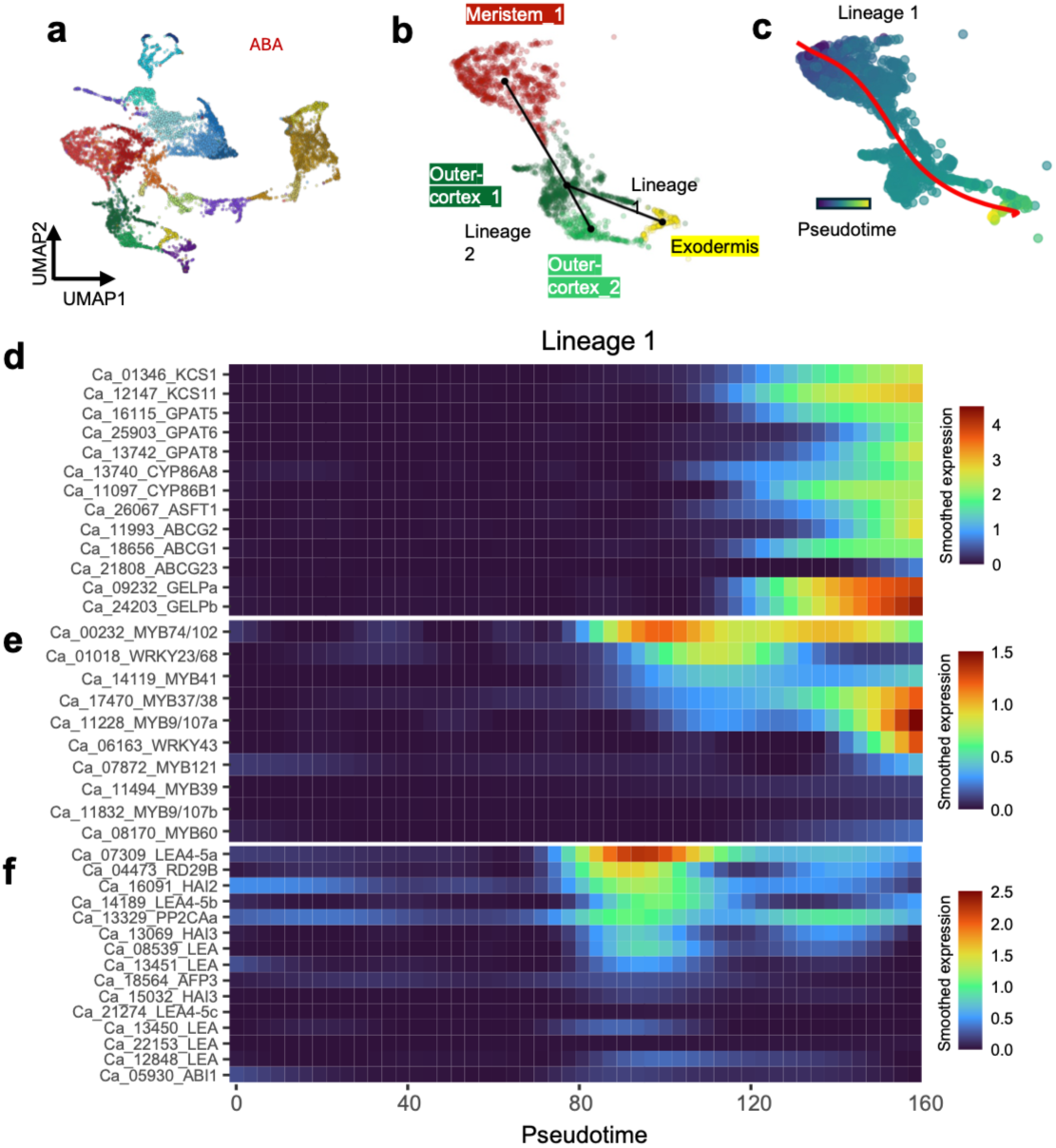
Pseudotime estimation recapitulates the temporal dynamics of *MYB* and ***WRKY* leading the differentiation of exodermis in chickpea roots. a,** Cell lineage detection using the Slingshot package (Street et al. 2018) revealed two lineages in the outer ground tissue (outer cortex and exodermis). The Meristem_1 cluster was included to help anchor the cell lineage detection and pseudotime estimations. Lineage 1 consists of clusters Meristem_1, Outercortex_1 and Exodermis_1, while Lineage 2 consists of Meristem_1, Outercortex_1 and Outercortex_2. **b**, Pseudotime trajectory through cell populations of Lineage 1. **d-f**, Smoothed expression profiles of suberin biosynthesis (**d**), *MYB* and *WRKYs* TFs induced by ABA in the bulk RNA-seq dataset (**e**) and canonical ABA-responsive genes (**f**) along the Lineage 1 pseudotime trajectory.

**Figure 7.**
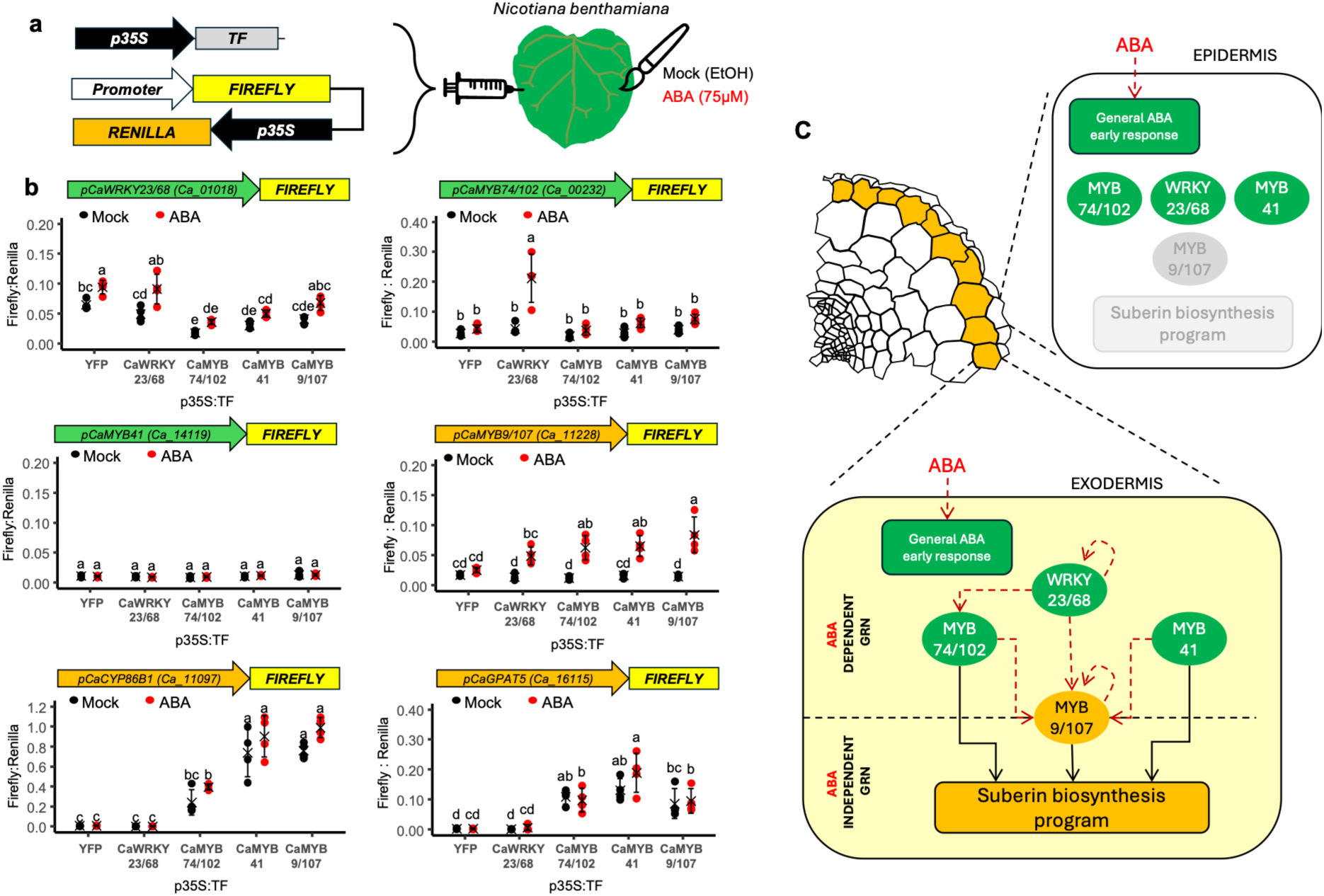
Transactivation assays of the ABA-dependent and ABA-independent gene regulatory networks controlling the suberin biosynthesis program in chickpea exodermis. **a**, Schematic diagram of transactivation assays in *Nicotiana benthamiana* leaves. *N. benthamiana* leaves were co-infiltrated with i) the reporter vector carrying the firefly luciferase under the control of the promoter of interest and the transformation expression control (*p35S:RENILLA*) and (ii) vectors overexpressing each candidate TF (*p35S:TF*). A vector overexpressing the YFP protein was used as a transactivation negative control. After 24 hours, infiltrated leaves were brushed with a solution containing either 75 µM ABA or a mock solution. After overnight incubation (16–18 h), leaf sections were collected for transactivation analysis. **b**, The firefly to renilla luciferase activity ratios of each promoter in response to the TF (x-axis) in mock (black) and ABA (red) conditions are presented. The black mark indicates the mean and error bars represent the standard deviation (*n* = 4). Different letters indicate statistically significant differences (one-way ANOVA followed by a post-hoc Tukey test, *P* ≤0.05). All transactivation assays were carried out twice (Supplementary Fig. S17). **c**, The working model for the suberization of chickpea exodermis. An ABA dependent gene regulatory network (GRN) involves the activation of expression of early responsive TFs, CaWRKY23/68 and two downstream *MYBs*, *CaMYB74/102 and CaMYB9/107* in an ABA dependent manner. These *MYBs* and *CaMYB41*, but not WRKY23/68, drive the expression of the tested suberin biosynthesis genes in an ABA-independent manner. Green color indicates expression in early modules, yellow in late modules. Red arrows indicate a positive interaction in presence of ABA, and black arrows indicate positive interaction regardless of ABA treatment.

### ABA-dependent and ABA-independent gene regulatory networks controlling exodermis suberization

To test our hypothesis, we used a trans-activation assay in *Nicotiana benthamiana* leaves (Fig. 7a). In this assay, we tested three hypotheses. Firstly, we tested if an ABA treatment in this orthologous system was able to induce expression of our target genes (Fig. 7a). In addition to *CaMYB74*/*102* (*Ca_00232*) and *CaWRKY23/68* (*Ca_01018*) promoters, we also tested those of *CaMYB9/107* (*Ca_11228*), *CaMYB41* (*Ca_14119*) and two ABA-induced, exodermis-specific suberin biosynthesis genes (*CaCYP86B1 (Ca_11077*) and *CaGPAT5* (*Ca_16115*)). Interestingly, we observed a mild induction of only *pCaWRKY23/68* in response to ABA in the YFP control (Fig. 7b, Supplementary Fig. S17), which suggests that WRKY23/68 could act as a downstream target of canonical ABA signalling pathway. Secondly, we tested if other candidate TFs, CaMYB74/102, CaWRKY23/68, CaMYB41 or CaMYB9/107, could induce expression of each other in the presence or absence of ABA. CaWRKY23/68 induced the expression of *CaMYB74/102* and *CaMYB9/107*, but only in the presence of ABA application (Fig. 7b, Supplementary Fig. S17). This indicates a post-translational effect on CaWRKY23/68 activity by ABA signalling, either by a direct modification such as phosphorylation (Bakshi and Oelmüller 2014), or by presence of required cofactors. Additionally, expression from *pCaMYB9/107* was induced by all tested TFs, but again, only in the presence of ABA (Fig. 7b, Supplementary Fig. S17). This activation of *pCaMYB9/107* by multiple TFs in an ABA-dependent manner indicates it acts downstream from the other TFs which is in agreement with its late expression in both the time-course (Fig. 4d) and the pseudotime trajectory (Fig. 6e), but at odds with its exodermis-specific expression as two of the upstream TFs have wider patterns (Fig. 5i). In our profiled cells, only cells in the exodermis cell-cluster express both *CaMYB9/107* and its putative upstream TFs (Supplementary Fig. S18), so a permissive regulation and self-induction may be employed to amplify transient signals and enforce suberin biosynthesis in response to ABA (Fig. 7b, Supplementary Fig. S17). Finally, we tested if our candidate TFs can activate genes in the chickpea suberin biosynthesis pathway. All tested CaMYBs, but not the CaWRKY23/68, were able to drive expression from *pCaCYP86B1* and *pCaGPAT5* both in an ABA-independent manner (Fig. 7b, Supplementary Fig. S17). This allowed us to propose a gene regulatory network linking ABA to suberin biosynthesis via CaWRKY23/68 transcriptional and post-translational activation and the downstream activation of suberin-related and exodermis expressing MYB TFs (Fig. 7c).

## Discussion

Our characterization of ABA-induced deposition of cell wall polymers in galegoid legume cortical layers uncovered diversity of phenotypes representing diverse differentiation trajectories (Fig. 1, Supplementary Fig. S1). We resolved some previous unclarities of presence of exodermis, e.g. in pea (Taleisnik 1999; Bani et al. 2018), to be due to ABA-mediated, i.e. environmental, induction of suberin and lignin. For chickpea, we show that the differentiated exodermis containing SL (Fig. 2c) and non-polar lignin deposition (Supplementary Fig. S1) acts as a functional apoplastic barrier (Fig. 2d). Given the infrequent occurrence of exodermis in the legume clade, chickpea represents a novel model for this cell type that provides additional resilience traits against environmental stressors (Sharma et al. 2023; Cantó-Pastor et al. 2024).

We also observed one species, *V. sativa*, where clear FY and BF signal in multiple outermost cortical layers (Fig. 1, Supplementary Fig. S1) did not correlate with presence of SL (Fig. 2c) or apoplastic barrier to PI diffusion (Fig. 2d). While SL was not observed in *V. sativa*, we hypothesize that the compounds detected by the FY histochemical assay are related to suberin, possibly its monomers unable to polymerize. It may be that *V. sativa* represents a snapshot of suberin deposition pathway evolution, i.e. loss or gain of the complete set of enzymes and transporters needed to be expressed at the right place. Discovery of this intermediate phenotype between presence and absence of exodermis will provide opportunities for research of root barrier evolution.

Transcriptional investigation of the chickpea exodermis differentiation trajectory with time-course and single-cell approaches of exogenous ABA treatment revealed a group of WRKY and MYB TFs that putatively connect general ABA signalling with exodermis-specific suberization (Fig. 7c). Certain WRKY (Krishnamurthy et al. 2020, 2021; Jo et al. 2025) and MYB (Barberon et al. 2016; Lashbrooke et al. 2016; Cohen et al. 2020; Shukla et al. 2021; Xu et al. 2022; Cantó-Pastor et al. 2024; Baumgart et al. 2025) family TFs are known for their roles in regulating suberin deposition. Similarly to tomato (Kajala et al. 2021; Cantó-Pastor et al. 2024; Jo et al. 2025), we identified MYB TFs that induce suberin biosynthesis genes (Fig. 7b) expressed in the cluster representing exodermis and not the endodermis (Fig. 5i). However, the CaWRKY23/68 we identified as the top-tier transcriptional activator of *CaMYB74/102* and *CaMYB9/107 (*Fig. 7b, c) plays a different role than the previously identified suberin-regulating WRKYs. In Arabidopsis, AtWRKY9 and AtWRKY33 directly bind the promoters of *CYP* genes of the suberin biosynthetic pathway (Krishnamurthy et al. 2020, 2021), while we did not observe CaWRKY23/68 activating the ABA-induced exodermis-specific *CaCYP8B1 (Ca_11097)*. In tomato, SlWRKY71 works antagonistically to suberin-inducing MYBs (Jo et al. 2025), repressing the expression of suberin biosynthesis genes. We observed no effect of CaWRKY23/68 onto the two suberin biosynthesis promoters (Fig. 7b) but detected a small repression of *pCaWRKY23/68* promoter activity by the suberin-inducing MYBs (Fig. 7b). This observation indicates that there is negative feedback between WRKYs and MYBs in chickpea (Fig. 7b) and tomato (Jo et al. 2025), even though working at different levels in the gene regulatory network.

Although ABA is widely known as a suberin-inducing signal acting via induction of the suberin-regulating MYBs (Kosma et al. 2014; Barberon et al. 2016; Shukla et al. 2021; Lu et al. 2022; Cantó-Pastor et al. 2024), the molecular mechanism how ABA triggers their induction has remained elusive. Our single-cell approach revealed that the canonical ABA signalling is activated broadly across many root cell types, showing a widespread ability of root cells to transcriptionally respond to ABA (Fig. 5e, f). However, the downstream transcriptional response driving suberin biosynthesis remains spatially restricted to the exodermis (Fig. 5g, h). This points to an exodermis-specific regulatory mechanism that can translate the canonical ABA signalling into the activation of the suberin gene-regulatory network. Here we provide a potential model that links ABA-dependent and independent gene regulatory networks in the chickpea exodermis (Fig. 7c). We observed that the TF activity of our top-tier TF, CaWRKY23/68, was induced by ABA in the transactivation assay (Fig. 7b), which could serve as such a mechanism. The activation is likely a post-translational effect as the assayed TFs were expressed under a strong near-constitutive promoter. WRKY TF activity is commonly strengthened by phosphorylation in Arabidopsis (Adachi et al. 2015; Zhou et al. 2024), making phosphorylation a good candidate for an induction mechanism. Furthermore, rice WRKY72 has been shown to be phosphorylated in response to ABA by the canonical ABA signalling component SnRK2 (SNF1-related protein kinase 2) (Park et al. 2009; Hou et al. 2019). It remains to be queried if WRKYs and their phosphorylation is the bridge between ABA perception and the canonical regulation of suberin by MYBs in various model systems.

It is not known how exodermis cell identity is positionally specified in any species, and similarly, our data did not allow identification of the cue that restricts the expression of *CaMYB9/107* and suberin biosynthesis genes specifically to the exodermis cluster (Fig. 5h). Additionally, expression of the early TFs, *CaWRKY23/68, CaMYB74/102* and *CaMYB41,* had wider domains, i.e. they were also expressed in clusters representing cell types overlaying the exodermis (root cap, trichoblast, Fig. 5h, Supplementary Fig. S14) in the presence and absence of ABA, which indicates restriction of the expression pattern along the progression of the network hierarchy, potentially through the expression of CaMYB9/107 and suggesting that the spatial or exodermis-identity cue does not necessarily act at the top level of this network module. ABA is made in response to environmental stresses, such as drought or salinity, and recently, it was shown that in rice, compaction-responsive ABA originates from internal tissues such as phloem (Zhu et al. 2025). Hence, the expression domains may be affected by the unusual direction or concentration of ABA, and further research would be needed to resolve if the wide expression patterns we observe for the early TFs (Fig. 5i) would be also observed in endogenous ABA-inducing abiotic stresses. Overall, our data provide evidence of functional sufficiency of the WRKY/MYB transcriptional module in the activation of the suberin biosynthesis program, expressed in chickpea exodermis in response to ABA treatment. How the expression pattern of this module is restricted to exodermis remains an exciting avenue for future research.

In addition to suberin gene expression unit in chickpea exodermis, there is also a lignin unit in response to ABA. We observed lignin biosynthesis genes to be expressed temporally and spatially in response to ABA to promote the lignification of the chickpea root exodermis (Supplementary Fig. S11c, d). Our data did not support clear candidates as regulators of lignin biosynthesis in exodermis (Supplementary Fig. S11e), but similar questions remain as for regulation of suberin biosynthesis. What sets exodermis apart from the other cell types in its gene regulatory capacity to induce lignin biosynthesis? What are the ABA-induced changes, possibly post-translational, that form the exodermis-specific, ABA-induced lignin biosynthesis unit?

All in all, we characterized the barrier occurrence and characteristics in several galegoid legumes, providing the first comparative framework for understanding root barrier diversity and evolution in this clade. Our characterization of temporal and spatial gene expression patterns in chickpea in response to exogenous ABA reveals key regulatory modules associated with hormone perception and cell-type specific suberin deposition in the legume family. Looking ahead, comparative transcriptomic analyses across the galegoid family will be crucial for uncovering the evolutionary innovations underlying the phenotypic diversity we observed. The conservation and divergence of temporal and spatial gene expression patterns (Fig. 3d, 5, 6) and the regulatory interactions observed in chickpea (Fig. 7b, c) remains to be tested in non-exodermal galegoid legumes, and thus, will provide important insights into how legume root barriers have evolved.

## Author contributions

KK: Conceptualization, Supervision, Project Administration, Funding Acquisition.

LJ, RK, SB, MR, JH, VT, JB, HC: Investigation.

RBF, PD: Resources.

LJ: Data Curation, Formal Analysis.

LJ, RK: Visualization.

LJ, KK: Original Draft.

All authors: Writing – Review & Editing.

## Acknowledgements

We would like to thank Judith Boursin for pea seeds, Wouter Kohlen for *Medicago truncatula* seeds and extensive conversations, Tonni Andersen for *Lotus japonicus* seeds and support, Swati Mahiwal for HCR-FISH training, Rene Geurts for ARqua1 strain, Chenxin Li for sharing the protocol for the protoplast density gradient, and Laura Ragni and Bruno Guillotin for critical reading of the manuscript. We also thank Xiren Cao, Henri Essombé Kuoh, Menno Druif and Han Atlamaz for technical assistance, and the WUR TEM facility for imaging.

## Supplementary Material

Supplementary Table S1: Abundance of suberin monomers in legume roots in response to ABA

Supplementary Table S2: Gene expression and differentially expressed genes (DEGs) analysis in chickpea roots in response to ABA

Supplementary Table S3: Single-cell gene markers in chickpea roots in response to ABA Supplementary Table S4: Primers used for cloning

Supplementary Table S5: DEGs in protoplasting experiment Supplementary Figure S1. Fluorol Yellow staining of root sections.

Supplementary Figure S2. Quantification summary of Basic Fuchsin (BF) dye intensity across multiple cell layers of legume roots in response to ABA.

Supplementary Figure S3. Suberin deposition is commonly under developmental and environmental control as shown by its deposition along root developmental zones.

Supplementary Figure S4. ABA effect on the root growth of galegoid legumes.

Supplementary Figure S5. Biochemical and structural evaluation of ABA-induced cell wall modifications.

Supplementary Figure S6. Expression profiles of putative ABA response genes, suberin biosynthesis genes and lignin biosynthesis genes.

Supplementary Figure S7. Chickpea roots deposit histochemically detectable suberin and lignin in exodermis within 24 h.

Supplementary Figure S8. DNA motif enrichment in the module I-VI gene promoters.

Supplementary Figure S9. CaMYBs and CaWRKYs do not share one-on-one orthology with Arabidopsis TFs.

Supplementary Figure S10. Extended information of the single-cell atlas of chickpea root responses to exogenous ABA.

Supplementary Figure S11. Annotation of the endodermis cluster based on Casparian strip genes and expression of lignin biosynthesis and putative lignin regulators in the chickpea single cell atlas.

Supplementary Figure S12. Similarities of transcriptional responses to ABA in the bulk and single-cell transcriptomic data.

Supplementary Figure S13. Reproducibility of single-cell chickpea root atlas in response to ABA.

Supplementary Figure S14. Validation of exodermis cluster annotation with HCR-FISH and promoter fusions.

Supplementary Figure S15. Evaluating the transcriptomics changes in chickpea roots by cell-wall digestion during protoplast isolation.

Supplementary Figure S16. Pseudotime trajectory of Lineage 2 does not involve suberin genes.

Supplementary Figure S17. Replicate experiments of the transactivation assays.

Supplementary Figure S18. Candidate suberin-regulating TFs are often not expressed in the same cells.

## Data availability statement

All sequencing data and coding scripts will be openly released upon acceptance of the manuscript for publication.

---

**Supplementary Figure S1.**
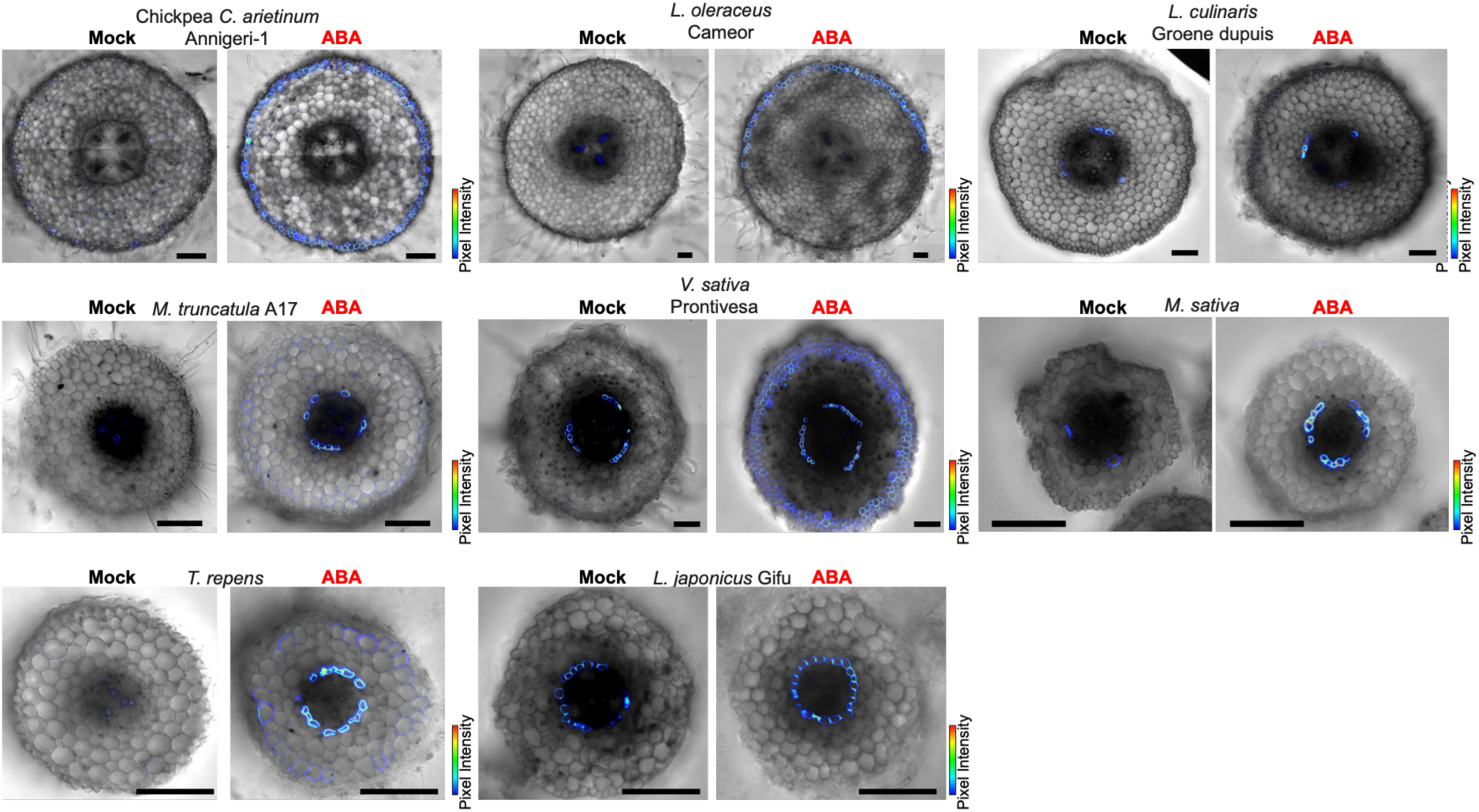
Fluorol Yellow staining of root sections. For each species, representative confocal images of full cross-sections of roots treated with mock or ABA are shown with the FY signal overlaid on top of bright field images (scale bars = 100 µm).

**Supplementary Figure S2.**
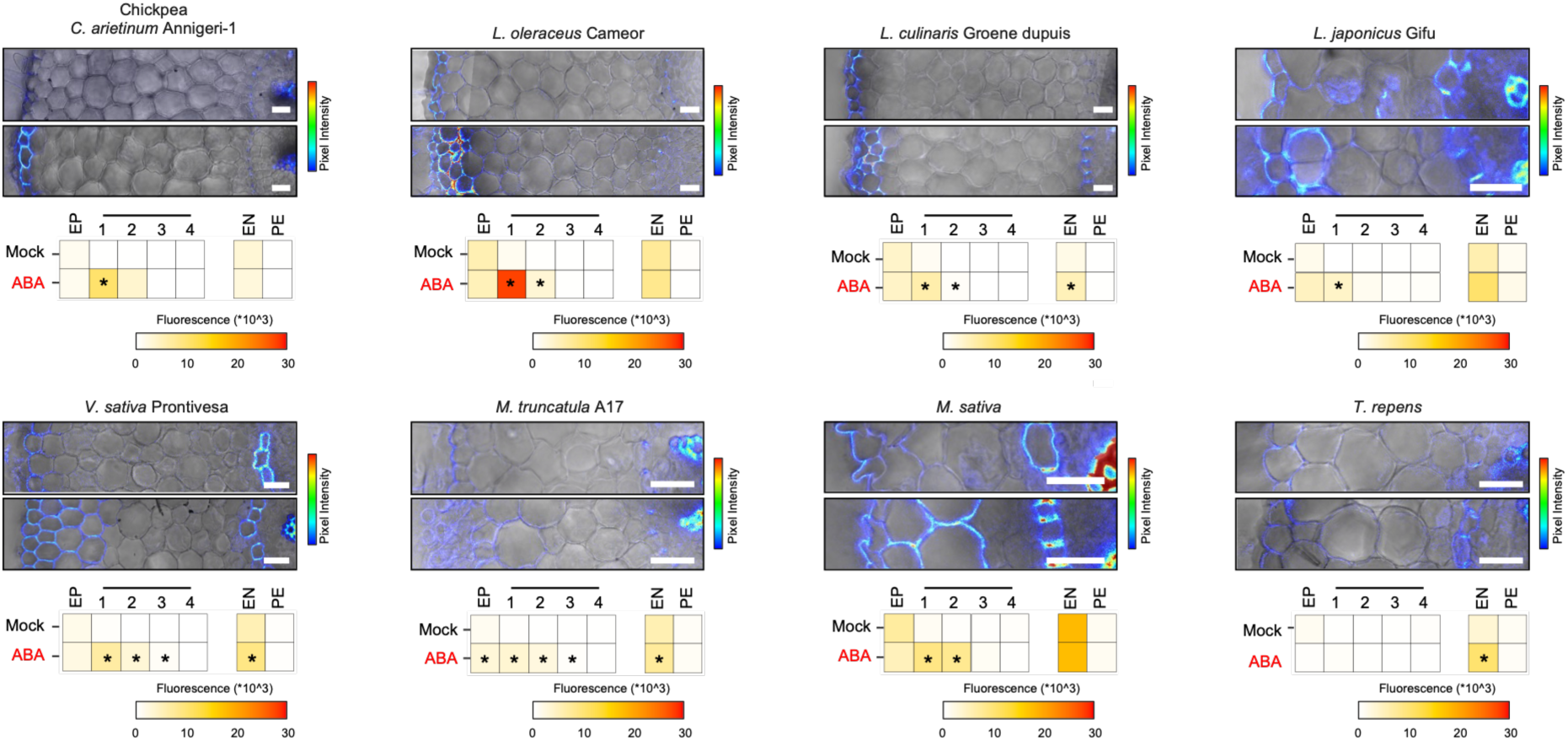
Quantification summary of Basic Fuchsin (BF) dye intensity across multiple cell layers of legume roots in response to ABA. For each species, the top images show representative confocal images of cross-section of roots treated with mock or ABA (2 µM) for 3 days, with the signal of BF overlaid on top of bright field images (scale = 25 µm). The bottom heatmap shows the average pixel intensity of the BF signal in different layers of the root cross section. Asterisks indicate significance based on Wilcoxon rank-sum test with a Benjamini-Hochberg adjusted P-value threshold of 0.05 was used to indicate the statistical significance (n ³ 8).

**Supplementary Figure S3.**
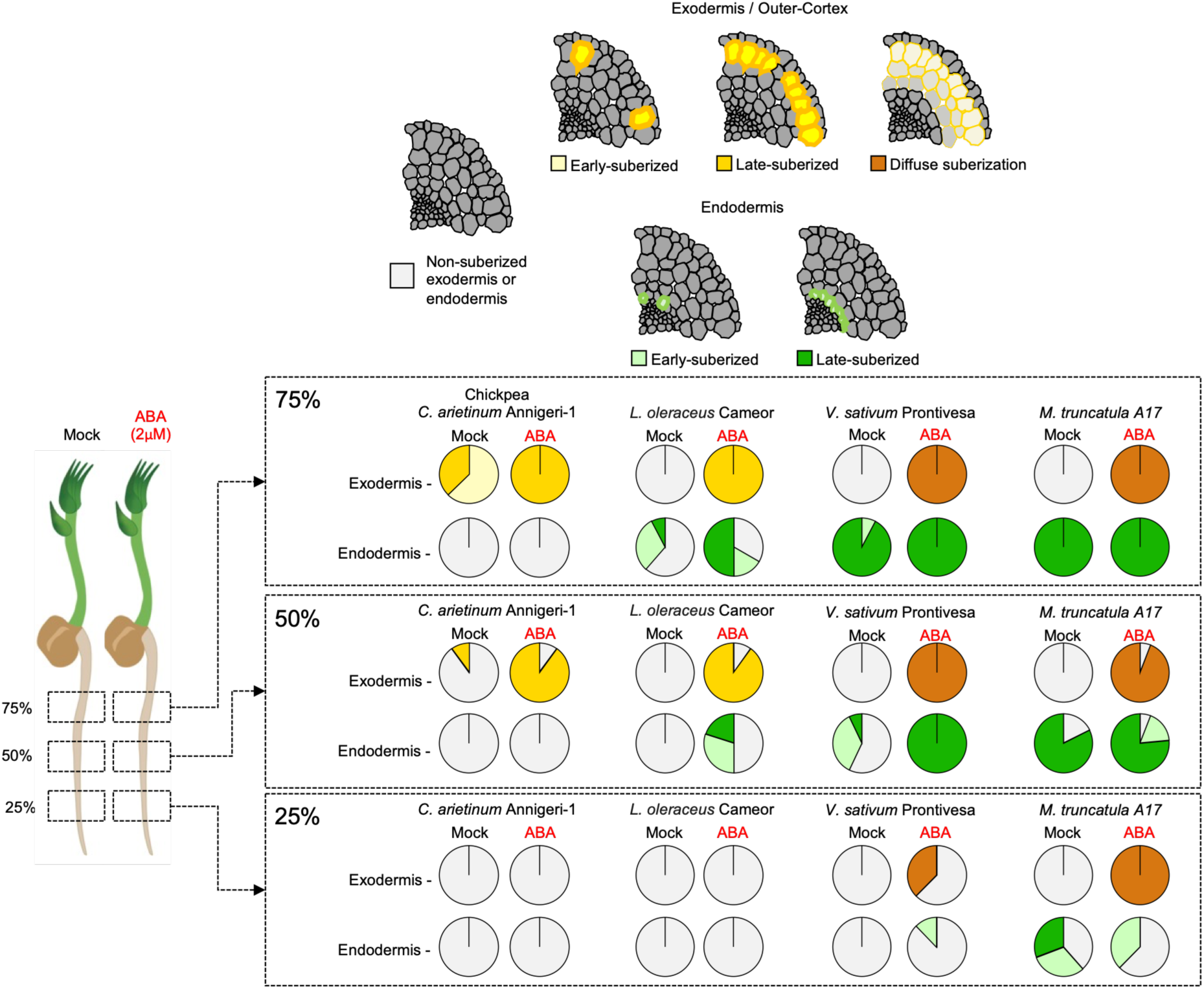
Suberin deposition is commonly under developmental and environmental control as shown by its deposition along root developmental zones. Suberin deposition in distinct developmental zones (25, 50 and 75% region of the root) in the exodermis/outer cortex or endodermis of chickpea (*C. arietinum*), pea (*L. oleraceus*), *V. sativa* and *M. truncatula* roots treated with mock or 2 µM ABA for 3 days. Pie charts depict the percentage of cross sections showing a non-suberized, an early-suberized (suberin in less than 50% of the cells in that layer), a late-suberized (suberin in more than 50% of cells in that layer), or diffuse (weaker than endodermal signal, multiple layers) suberization pattern in the cortex. n = 7-9 roots.

**Supplementary Figure S4.**
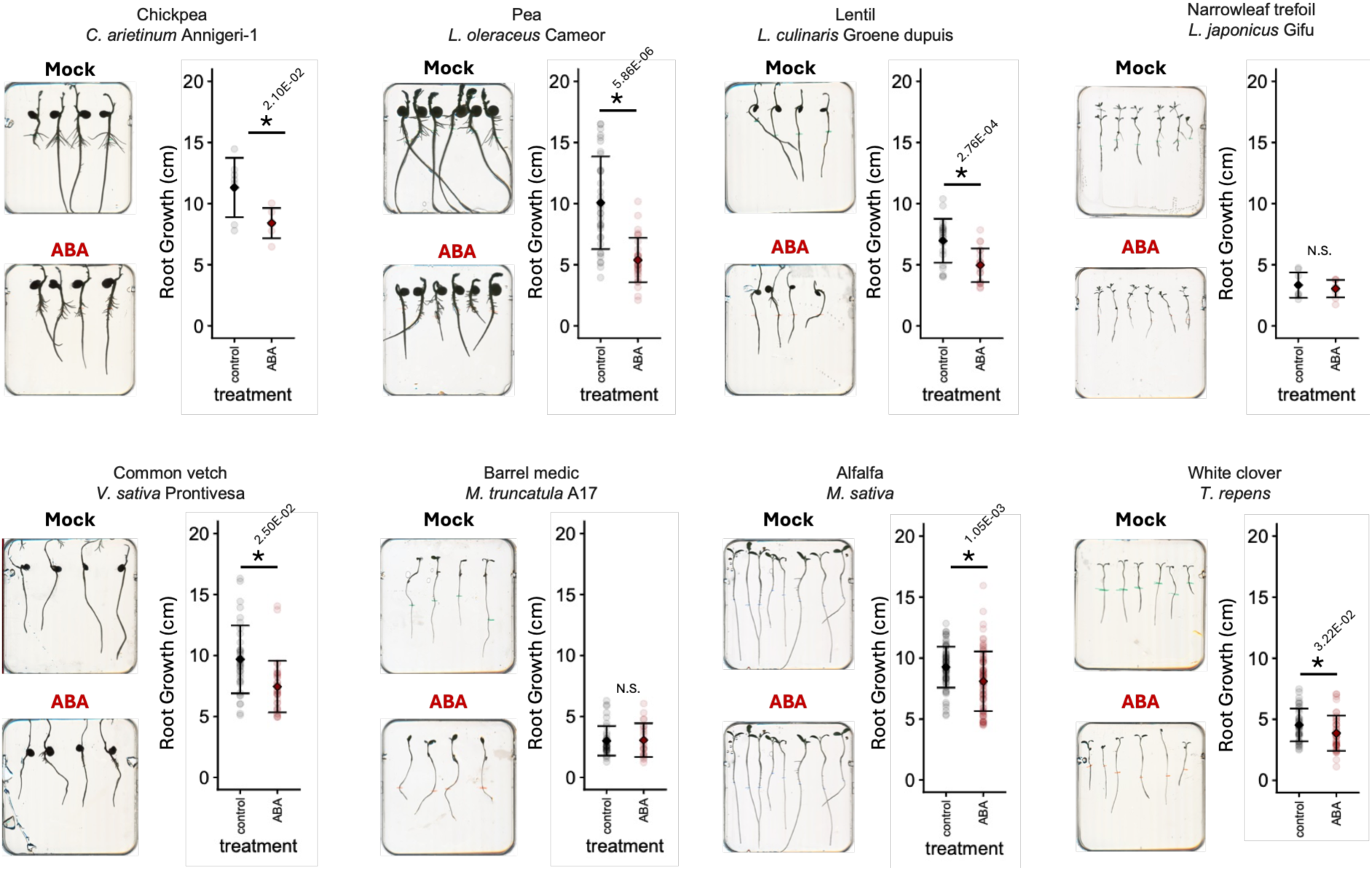
ABA effect on the root growth of galegoid legumes. ABA is known to inhibit root growth and change the lengths of developmental zones. Representative images of the root that were screened with FY (Fig. 1c) and BF (Supplementary Fig. S2) for each species and treatment, and the quantification of the root lengths. For all screened species except *M. truncatula and L. japonicus*, ABA inhibited root growth.

**Supplementary Figure S5.**
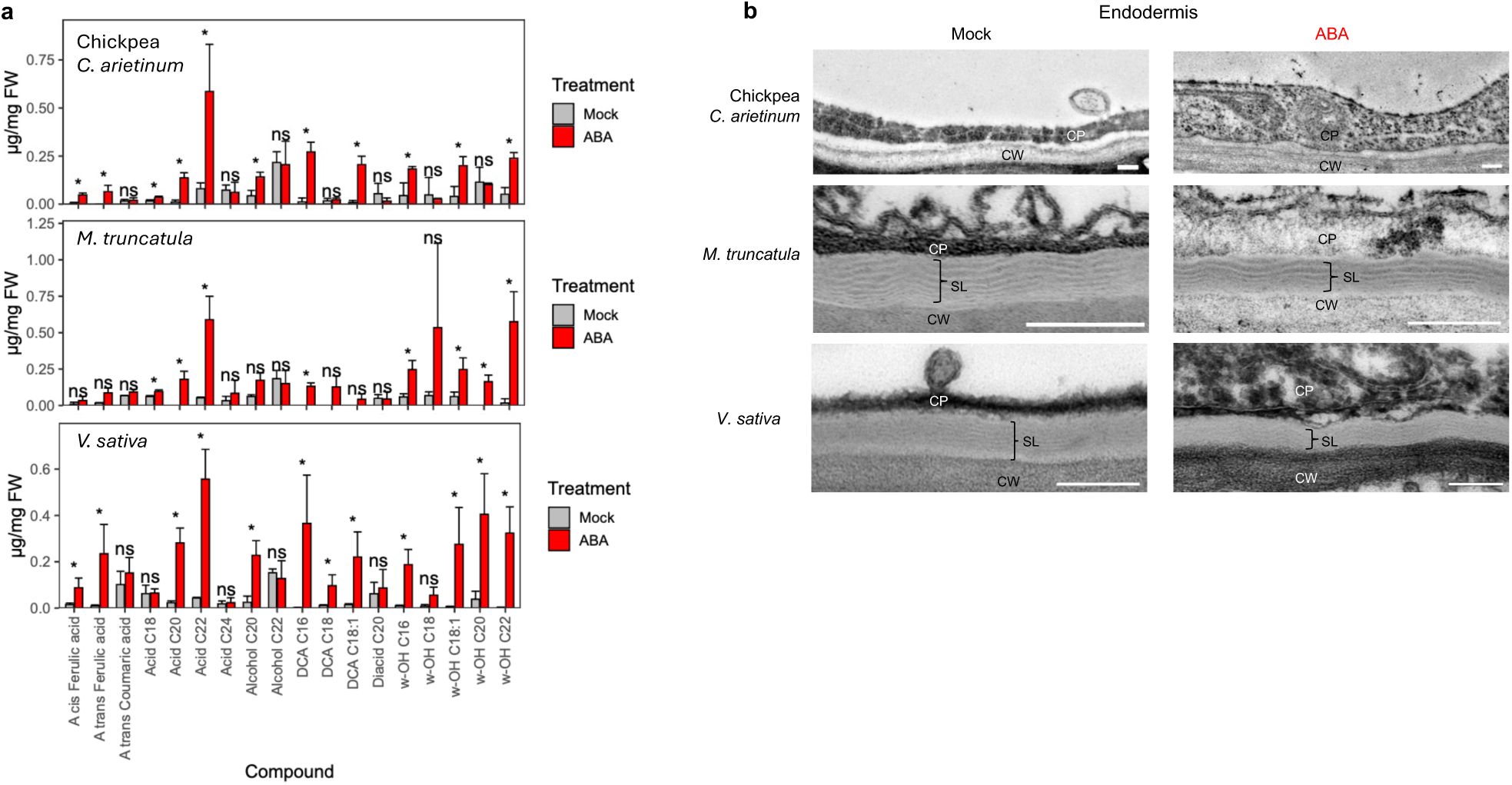
Biochemical and structural evaluation of ABA-induced cell wall modifications. **a**, Bar charts showing the abundance (µg/mg of fresh-weight (FW)) of individual suberin monomers in roots treated with mock (grey) or 2 µM ABA (red) for 24 hours. Asterisks indicate significance based on Student’s t-test with a P-value threshold of 0.05 (n = 4). **b,** Endodermis cell wall shown in TEM images of cross-sections at the 75% root region of Chickpea, *M. truncatula* and *V. sativa* treated with mock or 2 µM ABA for 3 days. The cell wall depicted in the images refers to the radial/anticlinal cell wall between two adjacent endodermal cells. Scale bar, 200 nm. SL: suberin lamellae; CW: cell wall; CP: cytoplasm.

**Supplementary Figure S6.**
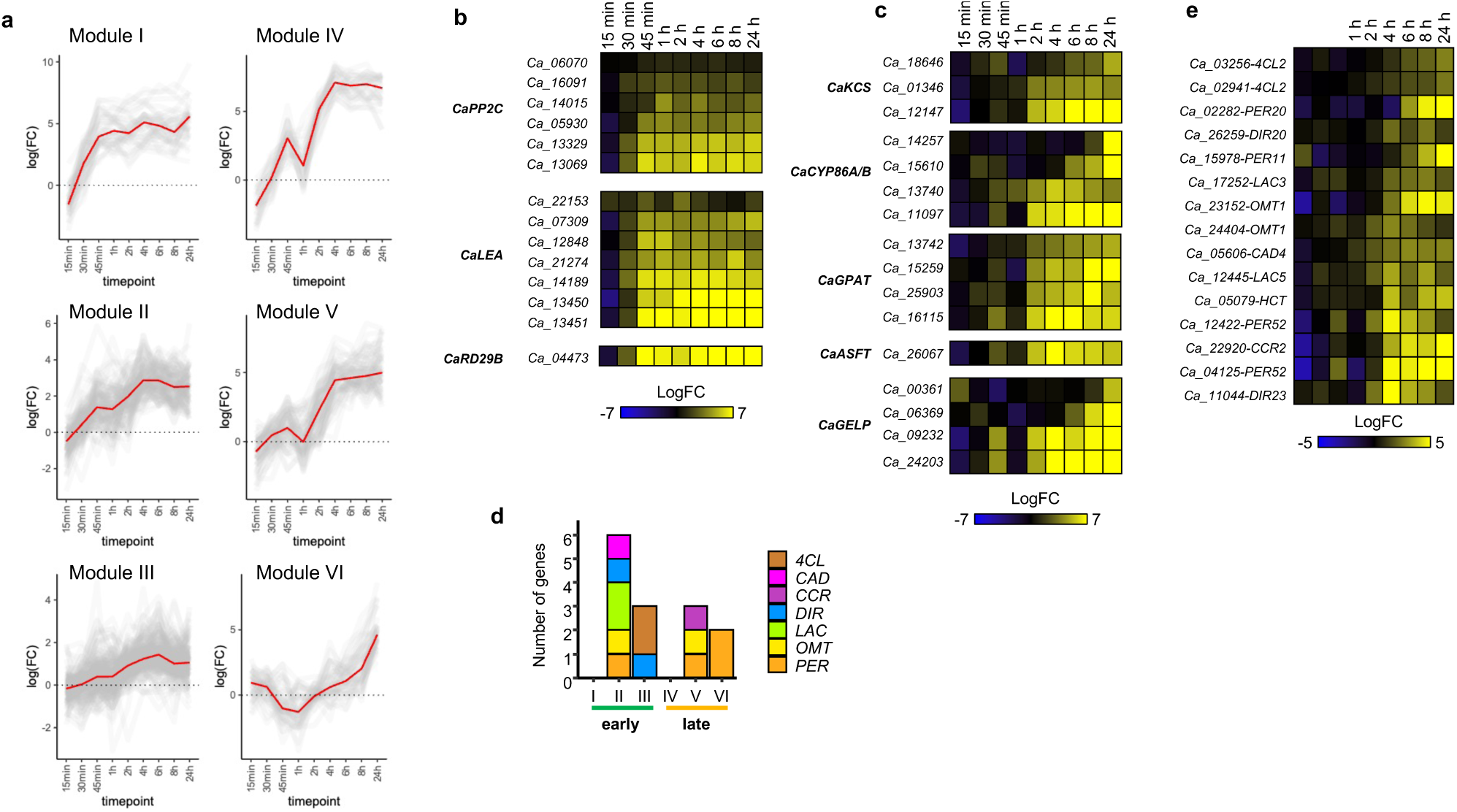
Expression profiles of putative ABA response genes, suberin biosynthesis genes and lignin biosynthesis genes. **a**, Lineplot showing the log of fold change (FC) of all genes (grey) in modules I-IV. Red lines depict median logFCs in each module. **b**, Heatmap showing the log (FC) of putative ABA response genes identified as DEGs. **c**, Heatmap showing the log (FC) of suberin biosynthesis genes identified as DEGs. **d**, Number of chickpea genes related to lignin biosynthesis identified in modules I-VI. **e**, Heatmap showing the log (FC) of lignin biosynthesis genes identified as DEGs.

**Supplementary Figure S7.**
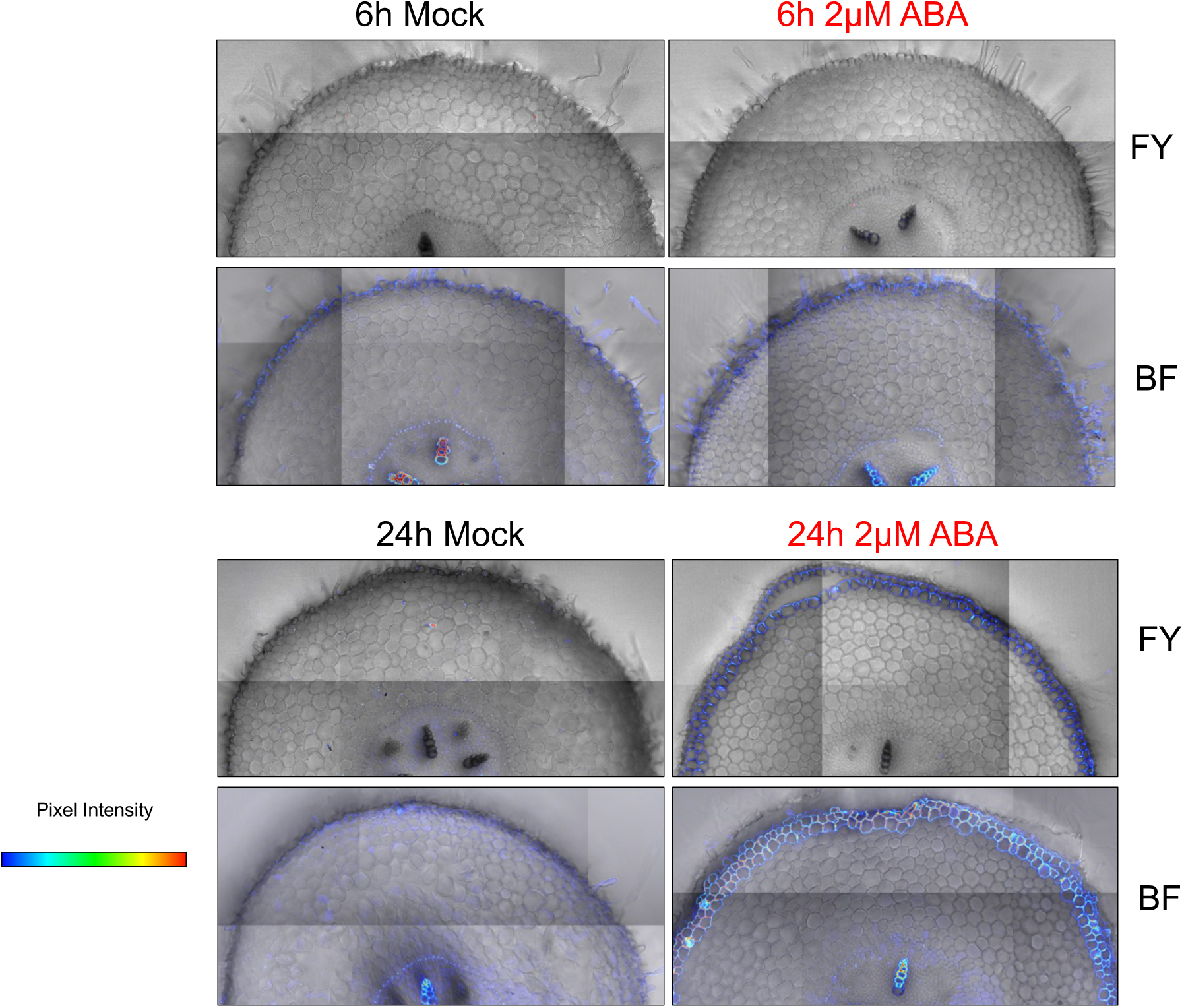
Chickpea roots deposit histochemically detectable suberin and lignin in exodermis within 24. **h.** Representative confocal images of chickpea roots used for the time course RNA-seq experiment stained with FY and BF are shown. Roots treated with mock or 2 µM ABA at two distinct time points (6 and 24 hours) were cross-sectioned at 75% point as the 50% point shown typically measured was included in the material harvested for RNA-seq. At this developmental zone, additional layer with FY and BF signal can be observed. FY and BF signals are overlaid on top of bright field images. Scale bar = 100 µm.

**Supplementary Figure S8.**
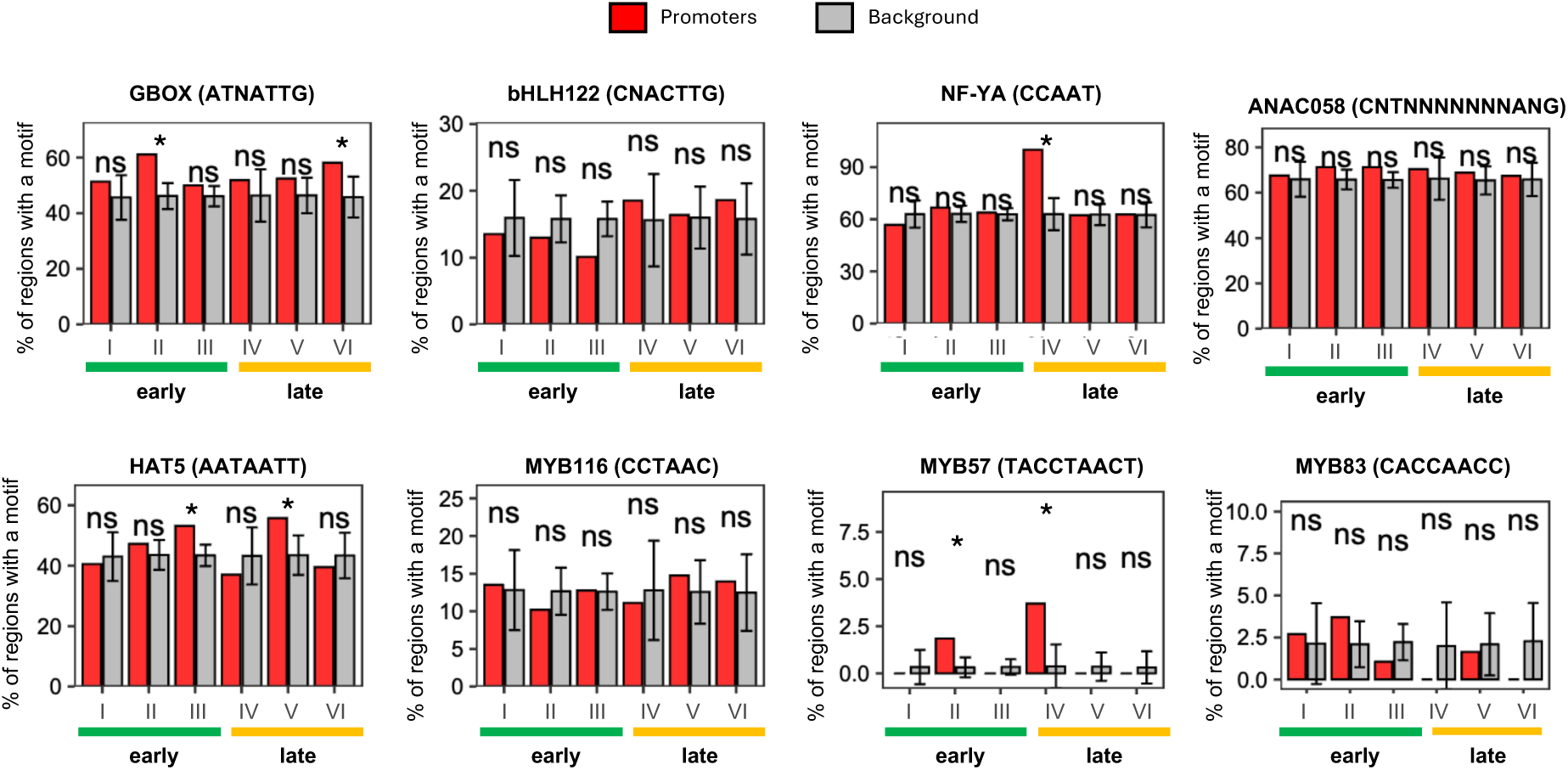
DNA motif enrichment in the module I-VI gene promoters. The selection of the above motifs was based on existing transcription factor families identified as DEGs (Supplementary Table S2). Bar graphs show the percentage of promoters of DEGs (red) and 1000 sets of promoters of randomly selected genes with equivalent number to the compared DEGs list (grey) containing the indicated DNA motifs. A Bonferroni-adjusted P-value threshold of 0.01 was used to indicate the statistical significance (*) of enrichment, while ns indicates no significance.

**Supplementary Figure S9.**
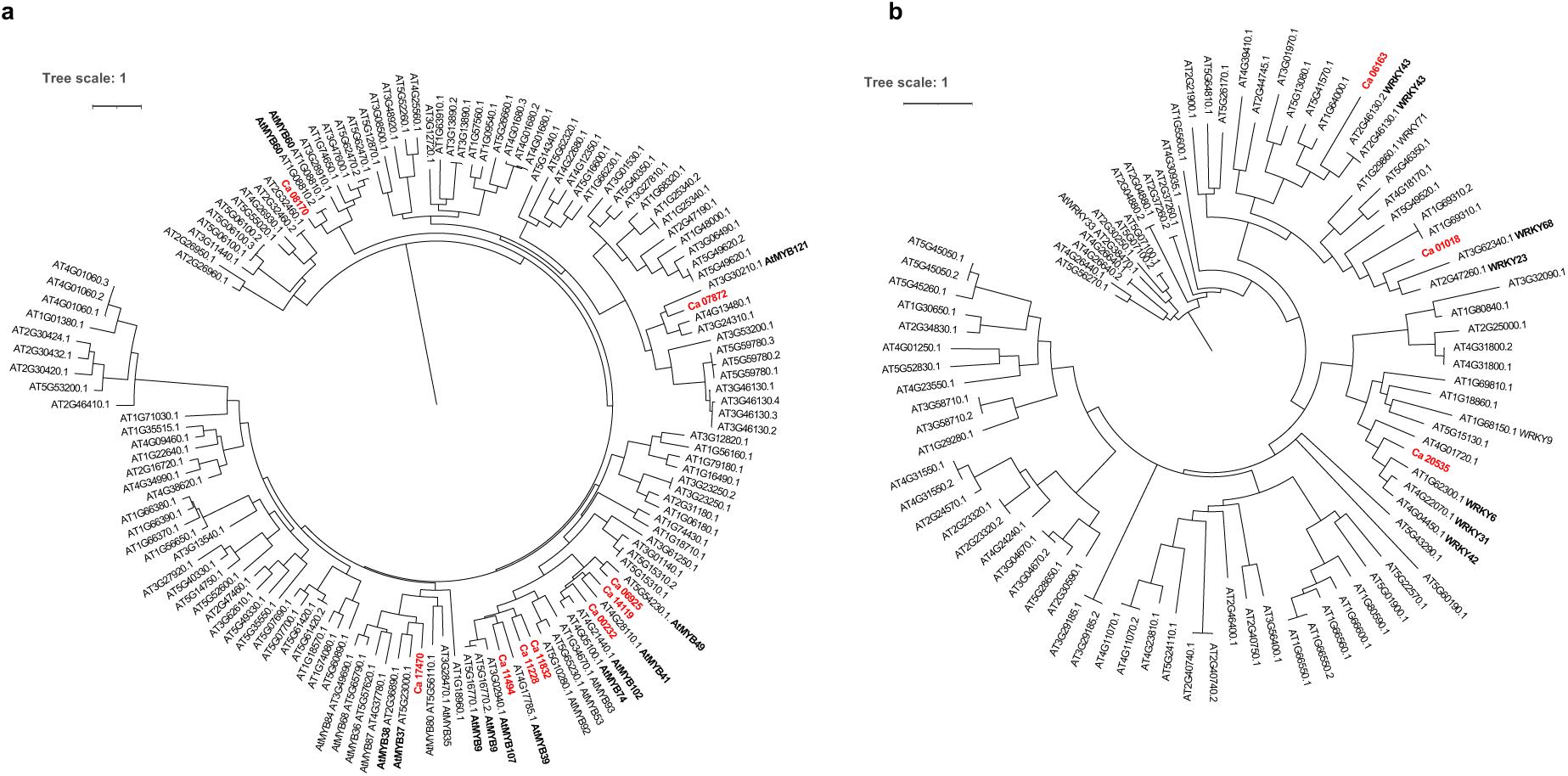
CaMYBs and CaWRKYs do not share one-on-one orthology with Arabidopsis TFs. Representative phylogenetic trees showing the closest Arabidopsis homologs to chickpea **a**, MYBs and **b**, WRKY TFs identified as upregulated by ABA in the time-course bulk RNA-seq experiment. Full length amino acid sequences of chickpea upregulated TFs and all annotated Arabidopsis MYB and WRKY TFs were aligned using MAFFT. The phylogenetic tree was determined using the maximum likelihood method with 1000 bootstraps using iQ-TREE. Distant nodes where chickpea MYB TFs were not detected were removed to improve the visualization of the cladogram. ABA-upregulated *CaMYBs* and *CaWRKYs* are indicated in red, and the TAIR10 symbol respective to close orthologues are indicated.

**Supplementary Figure S10.**
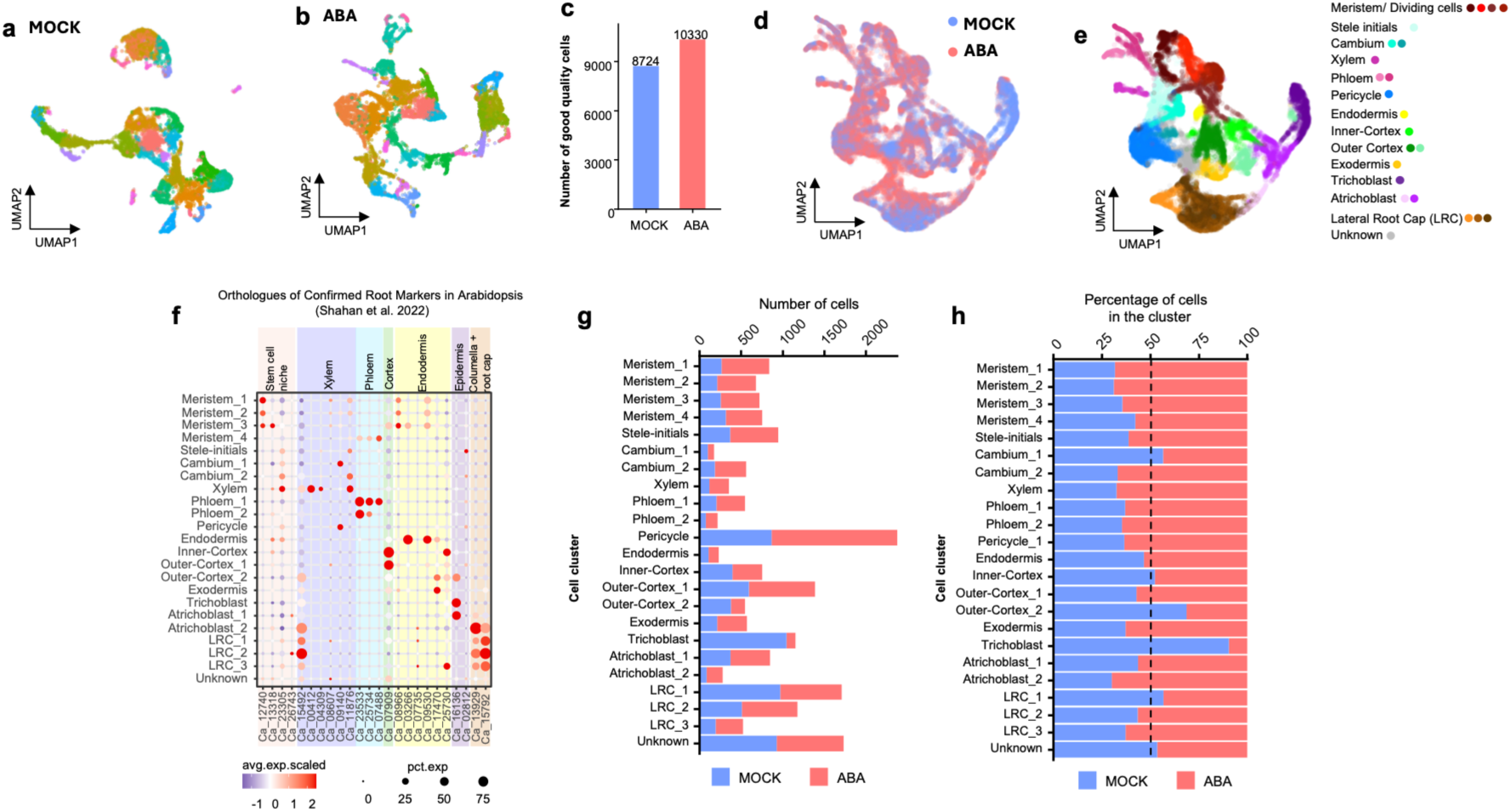
Extended information of the single-cell atlas of chickpea root responses to exogenous ABA. **a**, UMAP plot showing Seurat clusters of chickpea roots under mock conditions. **b**, UMAP plot showing Seurat clusters of chickpea roots treated with ABA. **c**, Number of high-quality cells in each dataset prior to integration. **d**, UMAP plot showing cell origin identities following data integration. **e**, Cell-type identities determined for each cluster. **f**, Dot plot showing the expression of chickpea orthologues corresponding to empirically determined root cell markers in Arabidopsis (Shahan et al., 2022). **g**, Number of cells detected in each cluster, categorized by original treatment: mock (lavender) and ABA (salmon). **h**, Proportion of cells from each treatment within each cluster.

**Supplementary Figure S11.**
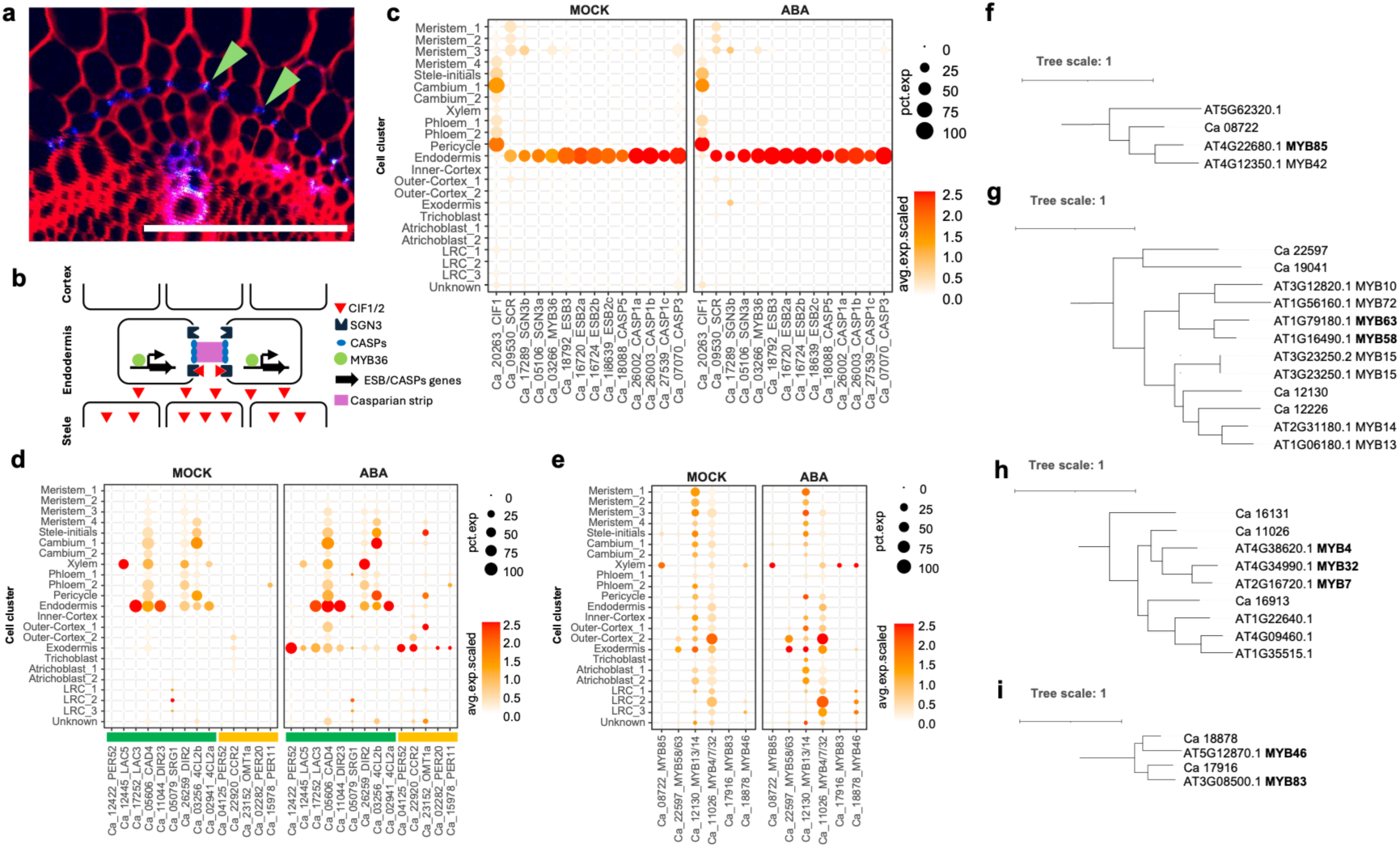
Annotation of the endodermis cluster based on Casparian strip genes and expression of lignin biosynthesis and putative lignin regulators in the chickpea single cell atlas. a, Cross section of chickpea roots stained with basic fuchsin (blue) and calcofluor-white (red) showing the localization of the lignified Casparian strip (green arrow) in the root endodermis. Scale bar, 100 µm. **b**, Conceptual diagram of CIF/SGN3 signaling pathway leading to Casparian strip formation. Adapted from (Fujita 2021). **c**, Dot plot showing the spatial expression in the single-cell data of genes related to the CIF/SGN3-mediated Casparian strip development. **d**, Dot plot showing the spatial expression of ABA-induced lignin biosynthesis genes. The color next to the gene name indicates their early (green) or late (yellow) temporal module in the bulk RNA-seq dataset. **e**, Dot plot showing the spatial expression in the single-cell data of homologs of known lignin-regulating *MYBs* in the single-cell dataset. **f, g, h, i,** Phylogenetic analysis identifying the relationships of closest chickpea homologs with the known Arabidopsis regulators of lignin biosynthesis: inducers AtMYB85 (**f**), AtMYB63 and AtMYB58 (**g**), inhibitors AtMYB4, AtMYB7 and AtMYB32 (**h**) and master regulators of secondary cell wall biosynthesis AtMYB46 and AtMYB83 (**i**).

**Supplementary Figure S12.**
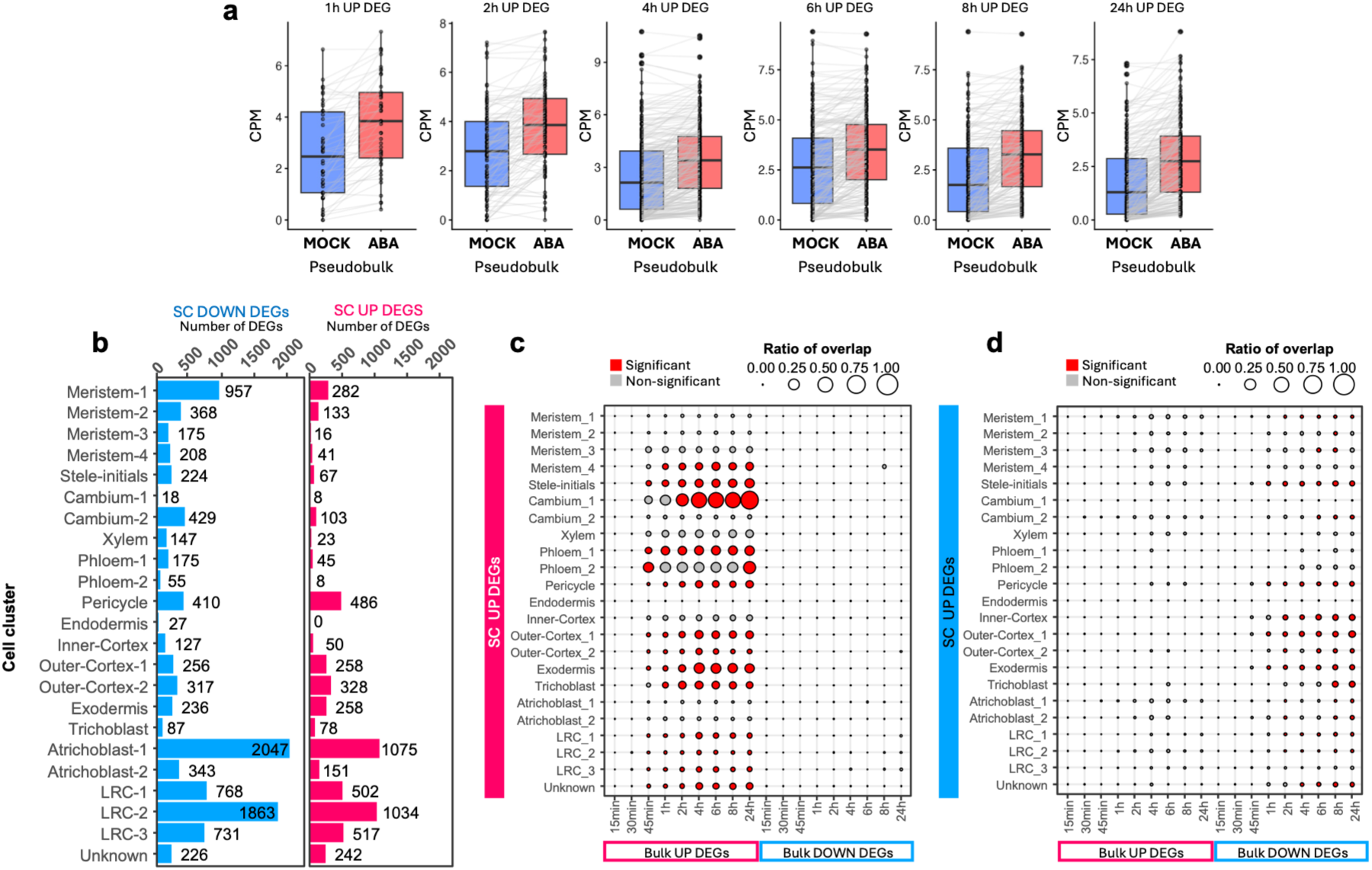
Similarities of transcriptional responses to ABA in the bulk and single-cell transcriptomic data. a, Pseudobulk expression profiles derived from the single-cell transcriptomics dataset revealed that genes previously characterized as ABA-upregulated via bulk RNA-seq exhibit a similar trend of expression. **b,** Number of differentially expressed genes (DEGs) upregulated (pink) and downregulated (blue) across cell clusters in ABA treated versus mock roots. **c**, Overlap between single-cell UP DEGs and temporal bulk RNA-seq UP and DOWN DEGs in the different timepoints. Circle size represents the overlap ratio between the two datasets, while color indicates the statistical significance of the overlap based on a hypergeometric test (P < 0.01). **d**, Overlap between single-cell DOWN DEGs and bulk RNA-seq UP and DOWN DEGs in the different timepoints.

**Supplementary Figure S13.**
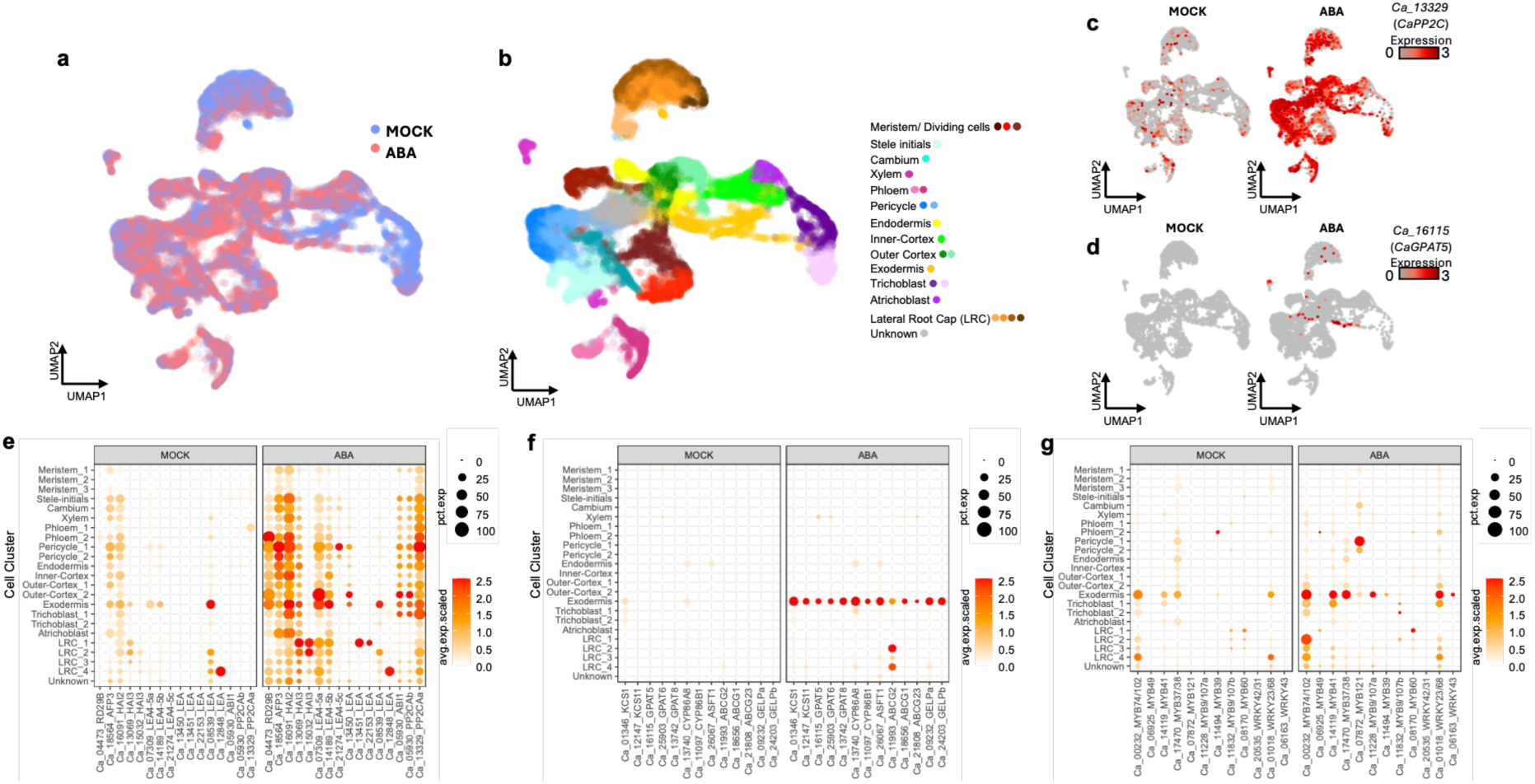
Reproducibility of single-cell chickpea root atlas in response to ABA. Second independent experiment of ABA-treated roots (48 hours) compared to the same mock control as Fig. 5 and Supplementary Fig. 10. A similar pattern is observed between experiments. Reproducibility as observed by **a,** UMAP. **b**, markers identified, and **c-g,** profile of expression of ABA responsive genes (**c,e**) and suberin biosynthesis genes (**d, f**) and MYB and WRKY TFs (**g**). Dot size indicates the percentage of cells in the cluster that expresses the specific gene (% expressed). Dot colors represent the average scaled expression of genes in the clusters (Avg. expression).

**Supplementary Figure S14.**
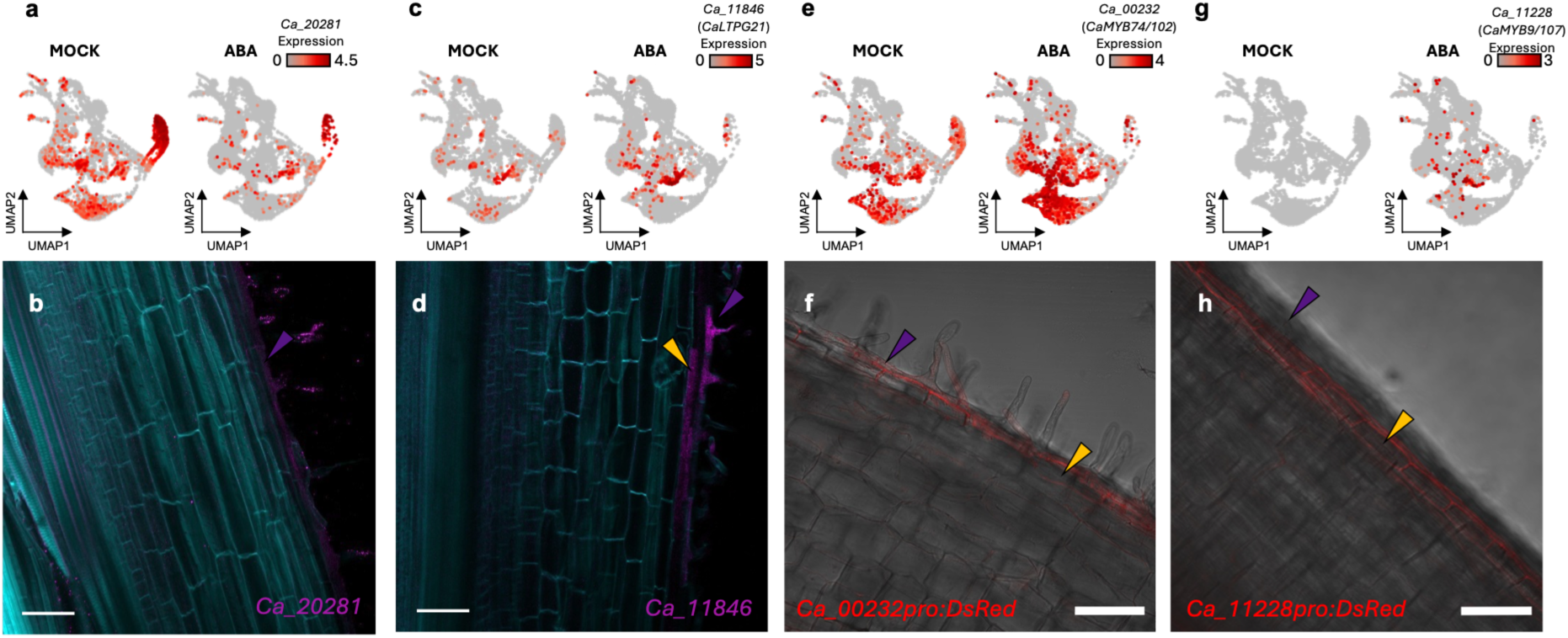
Validation of exodermis cluster annotation with HCR-FISH and promoter fusions. a, Transcript abundance of *Ca_20281* and **c,** *CaLTPG21 (Ca_11846)* in the chickpea mock and ABA single cell data and **b, d,** in chickpea roots treated with 2mM ABA for 24h visualized via HCR Gold RNA-FISH. Magenta represents the HCR Gold RNA-FISH probe fluorescence, cyan represents Calcofluor White. **e,** Transcript abundance of *CaMYB74/102* (*Ca_00232)* and **g,** *CaMYB9-107 (Ca_11228)* in the chickpea mock and ABA single cell data and **f, h,** in chickpea hairy roots transformed with *promoter:DsRed* reporter construct. Red represents DsRed fluorescence, overlaid with bright-field image of the root. Scale bars represent 100 mm. Yellow arrowheads point to exodermis (outermost cortex), purple arrowheads point to epidermis.

**Supplementary Figure S15.**
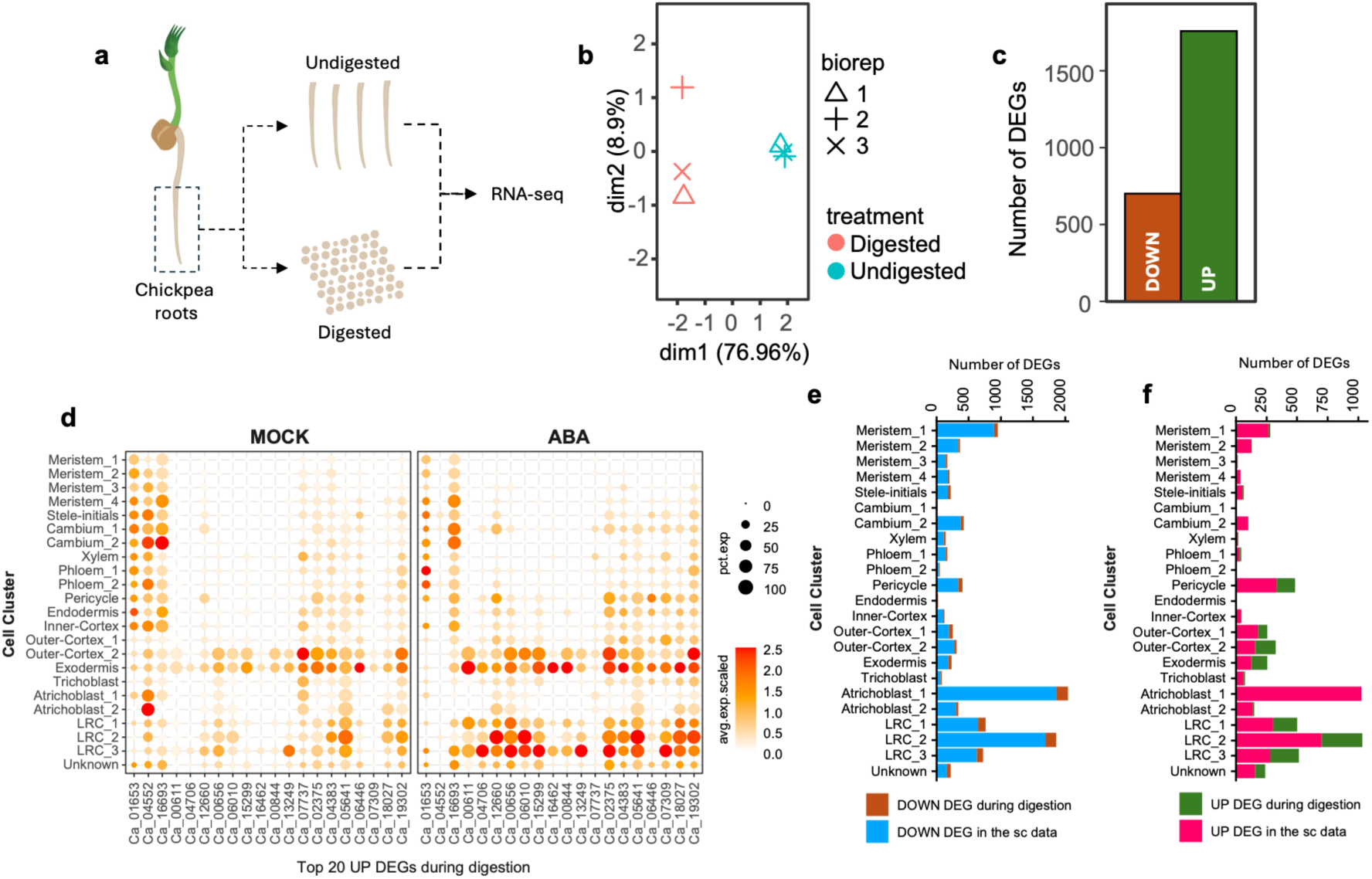
Evaluating the transcriptomics changes in chickpea roots by cell-wall digestion during protoplast isolation. **a**, Conceptual diagram of the experiment setting. Chickpea roots were protoplasted (digested) or not protoplasted (undigested), followed by transcriptome profiling through bulk RNA-seq. b, Multidimensional Scaling (MDS) plot showing the separation between digested and undigested RNA-seq samples. **c**, Number of differentially expressed genes (DEGs) identified from bulk RNA-seq data. Significance thresholds were set at an adjusted p-value of ≤ 0.01 and log2FC ≥ 1 for upregulated and log2FC ≤-1 for downregulated genes. **d,** Dot plot showing the expression of the top 20 UP DEGs in the digested roots, based on adjusted p-value, in the single-cell transcriptome data. The profile show that protoplast induced genes show relative similar expression profiles among mock and aba treated roots in the single cell data. **e**, Number of the single-cell DOWN DEGs that are down-regulated in the digested roots. **f**, Number of single-cell UP DEGs that are also up-regulated in the digested roots.

**Supplementary Figure S16.**
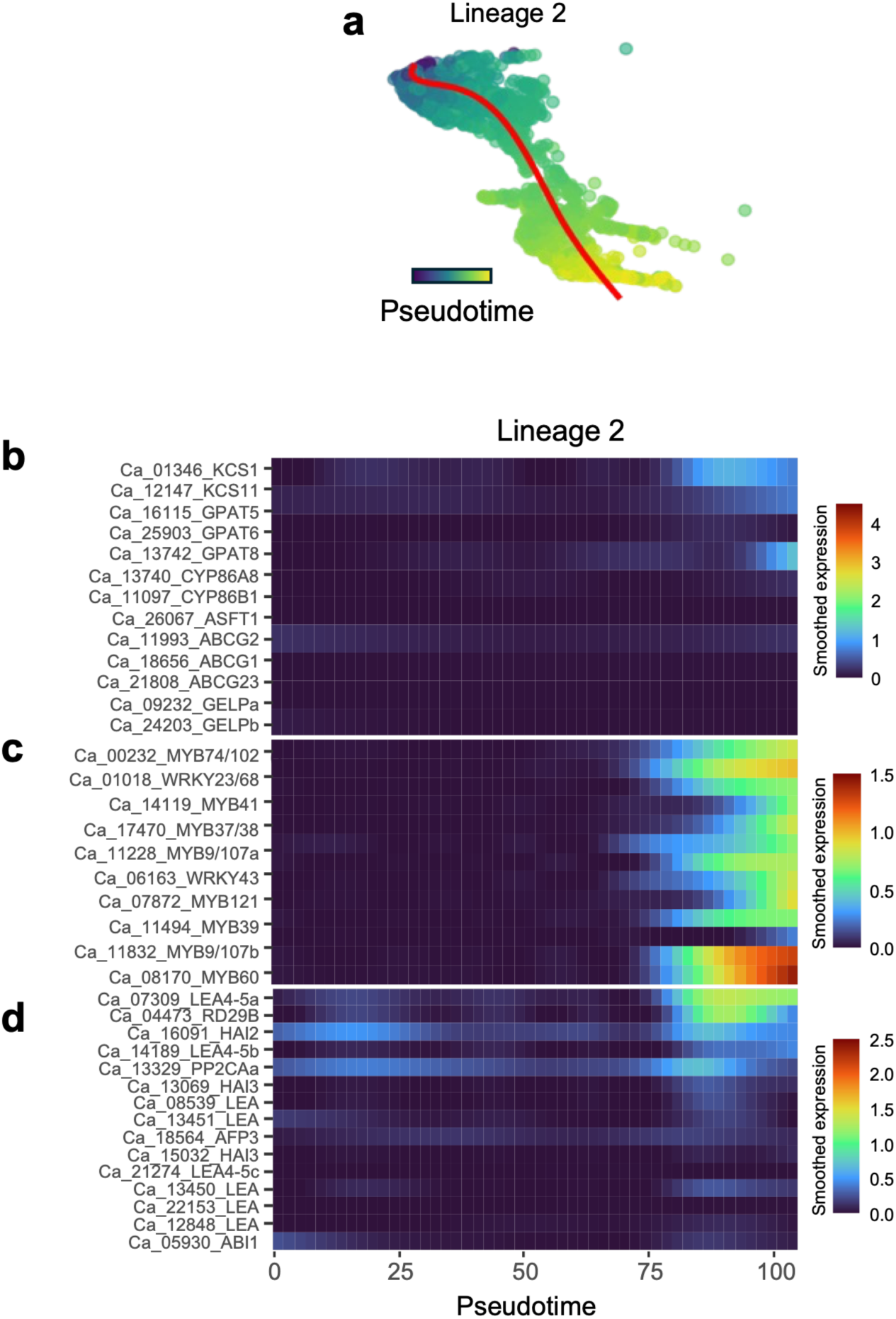
Pseudotime trajectory of Lineage 2 does not involve suberin genes. **a**, Pseudotime trajectory through cell populations of Lineage 2 (Meristem_1, Outercortex_1 and Outercortex_2). **b-d**, Smoothed scaled expression profiles of suberin biosynthesis (**b**), *MYB* and *WRKY* TF (**c**) and ABA-responsive genes (**d**) along Lineage 2 pseudotime trajectory

**Supplementary Figure S17.**
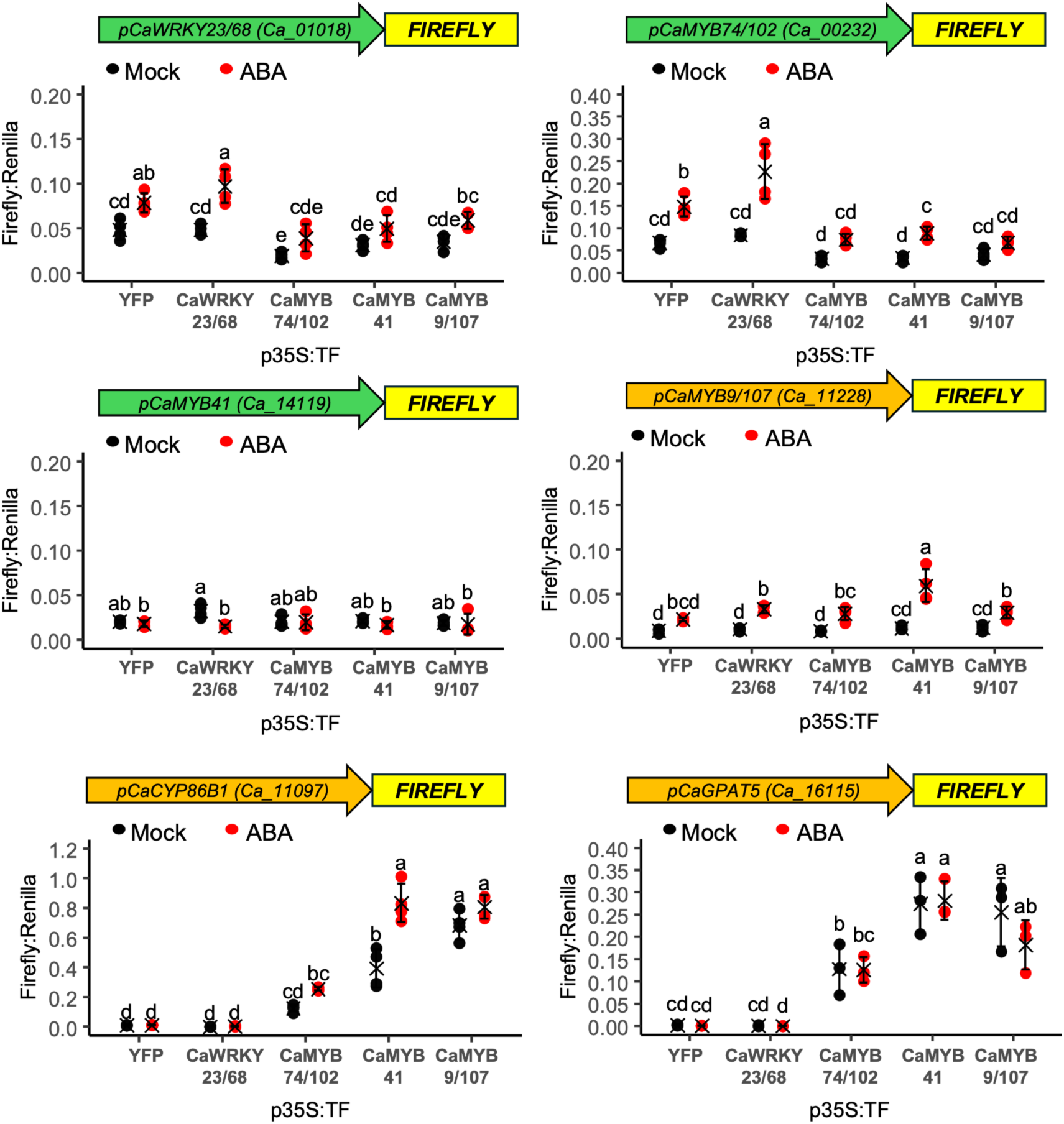
Replicate experiments of the transactivation assays. The transactivation assays (Fig. 7b) were replicated with similar results supporting the same conclusions.

**Supplementary Figure S18.**
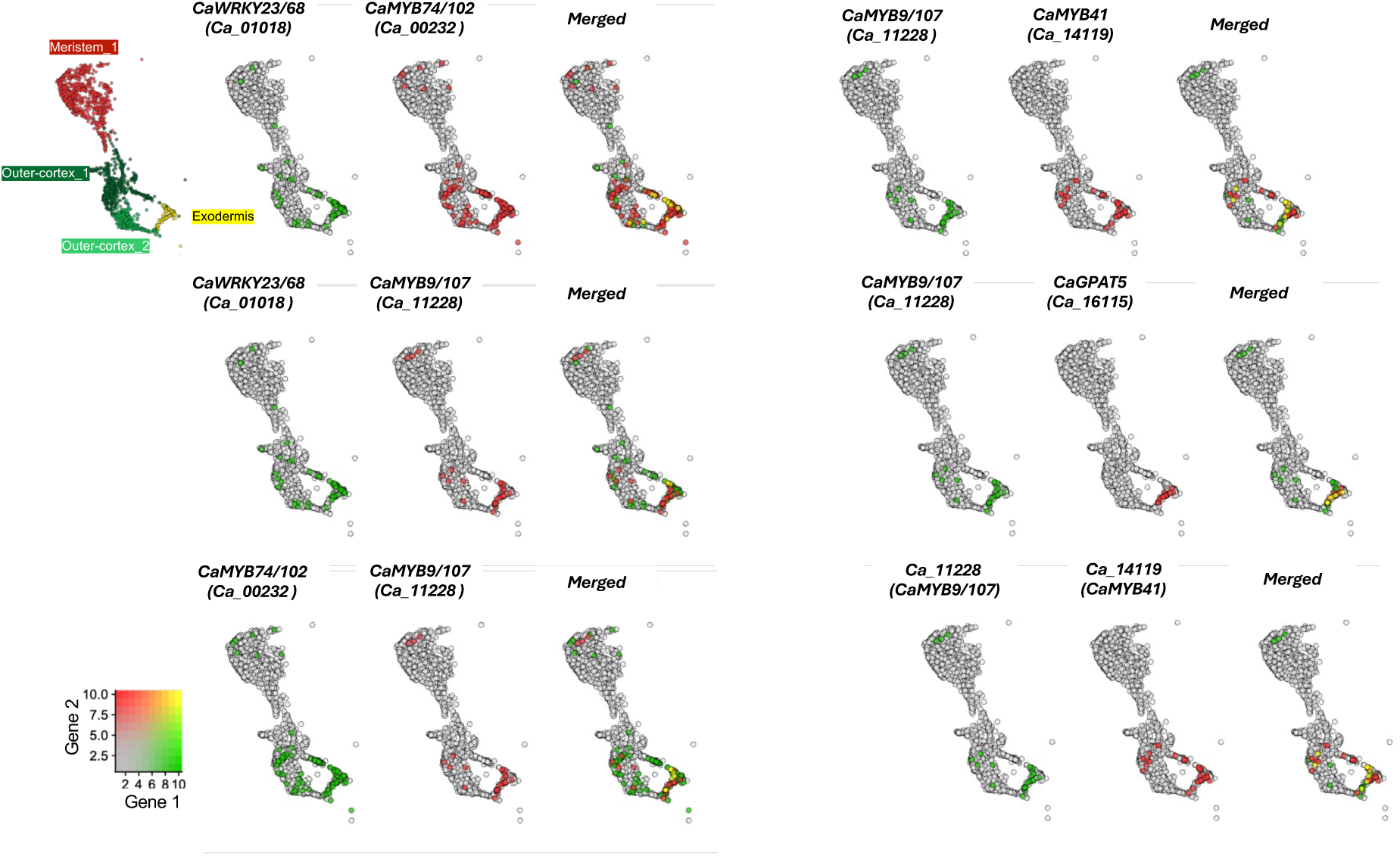
Candidate suberin-regulating TFs are often not expressed in the same cells. Overlap of expression in the single cell clusters representing outer cortex and exodermis for the genes encoding the candidate TFs (*CaWRKY23/68, CaMYB74/102, CaMYB9/107, CaMYB41)* and two suberin biosynthesis genes (*CaGPAT5, CaCYP86B1*). Gene 1 being expressed in a cell (blend threshold of 0.5) in green, gene 2 in red, and both expressed in the same cell in yellow. *CaMYB9/107* was induced by all candidate TFs (Fig. 7b) but it is often not expressed in the same cells as the other candidate TFs.

